# Generalized cue reactivity in dopamine neurons after opioids

**DOI:** 10.1101/2024.06.02.597025

**Authors:** Collin M. Lehmann, Nora E. Miller, Varun S. Nair, Kauê M. Costa, Geoffrey Schoenbaum, Khaled Moussawi

## Abstract

Cue reactivity is the maladaptive neurobiological and behavioral response upon exposure to drug cues and is a major driver of relapse. The leading hypothesis is that dopamine release by addictive drugs represents a persistently positive reward prediction error that causes runaway enhancement of dopamine responses to drug cues, leading to their pathological overvaluation compared to non-drug reward alternatives. However, this hypothesis has not been directly tested. Here we developed Pavlovian and operant procedures to measure firing responses, within the same dopamine neurons, to drug versus natural reward cues, which we found to be similarly enhanced compared to cues predicting natural rewards in drug-naïve controls. This enhancement was associated with increased behavioral reactivity to the drug cue, suggesting that dopamine release is still critical to cue reactivity, albeit not as previously hypothesized. These results challenge the prevailing hypothesis of cue reactivity, warranting new models of dopaminergic function in drug addiction, and provide critical insights into the neurobiology of cue reactivity with potential implications for relapse prevention.

---

Enduring relapse vulnerability remains a major challenge for the treatment of substance use disorders (Strang, Volkow et al. 2020) and attempts to curb relapse rates have not yielded significant improvements in the last fifty years (Sinha 2011). Exposure to drug-associated cues triggers craving and increases relapse risk(Saraiya, Jarnecke et al. 2021). This reflects the enhanced and enduring motivational effect of these cues (McHugh, Park et al. 2014). Such neurobiological and behavioral response is referred to as cue reactivity. Several models of addiction emphasize the role of cue reactivity and propose that addiction is a dopamine-dependent disorder of associative learning whereby repeated exposure to addictive drugs results in overvaluation and overpowering salience of drug cues through abnormally strong and long-lasting cue-drug associations (Di Chiara 1999, Berke and Hyman 2000, Redish 2004, Berridge and Robinson 2016, Kalhan, Redish et al. 2021).

Dopamine neurons provide a teaching signal that resembles reward prediction errors (RPEs) and modulates cue-reward associations (Schultz, Dayan et al. 1997, Steinberg, Keiflin et al. 2013, Eshel, Bukwich et al. 2015, Gershman and Uchida 2019). A leading hypothesis of cue reactivity is that addictive drugs, all of which increase dopamine signaling due to their direct pharmacological effects even when the cue-drug association is fully learned (Volkow, Fowler et al. 2004), result in enhanced maladaptive learning by causing a persistently positive RPE every time the drug is taken (Redish 2004, Keiflin and Janak 2015, Kalhan, Redish et al. 2021). Over time, this leads to a runaway enhancement of dopamine firing response to drug cues that approaches a maximum, resulting in overvaluation and triggering intense craving and relpase. Critically, this hypothesis posits a selective enhancement of dopamine response to drug cues, to the exclusion of other, non-drug cues. While this account is parsimonious, it has been subject to criticism (Panlilio, Thorndike et al. 2007, Marks, Kearns et al. 2010) and its underlying temporal difference RPE model of the dopamine signal has been criticized (supplementary note 1) (FitzGerald, Dolan et al. 2015, Gardner, Schoenbaum et al. 2018, Langdon, Sharpe et al. 2018, Gershman and Uchida 2019, Jeong, Taylor et al. 2022, Takahashi, Stalnaker et al. 2023). Further, the core prediction that dopamine RPE responses to drug-related cues should be selectively increased has not been tested. We conducted, across different institutions, two independent experiments using Pavlovian and operant procedures to directly test this prediction. We used single-unit recordings in rat ventral tegmental area (VTA) to compare dopaminergic firing responses (hereafter ‘dopamine response’) within the same neurons to cues associated with opioids (remifentanil) vs. natural rewards. Here we show that within-neuron responses are not different between cues predicting opioids vs. natural rewards in animals with a history of opioid exposure. Our data challenge the noncompensable RPE hypothesis of cue reactivity and offers a new understanding of how dopamine neurons respond to cues and rewards after repeated opioid exposure. Overall, these results shed new light into the role of dopamine in associative learning in drug addiction.

## Pavlovian procedure (Experiment 1)

We first validated a rat model that allows simultaneous single-unit recordings and intravenous (IV) delivery of the ultrashort-acting opioid remifentanil (RMF) (Fig. 1). RMF is a selective µ opioid agonist with similar reinforcing properties to other opioids (Panlilio and Schindler 2000) and a short half-life (0.3-1.1 min) (Crespo, Sturm et al. 2005). Rats (n = 11) were implanted with drivable microelectrode bundles targeting the VTA, and jugular IV catheters for IV fluid delivery. Rats were trained in a behavior box with a customized commutator to prevent tangling of the IV catheter line and headstage cable allowing for stable extended recording sessions (Fig. 1, a and b).

**Fig. 1.**
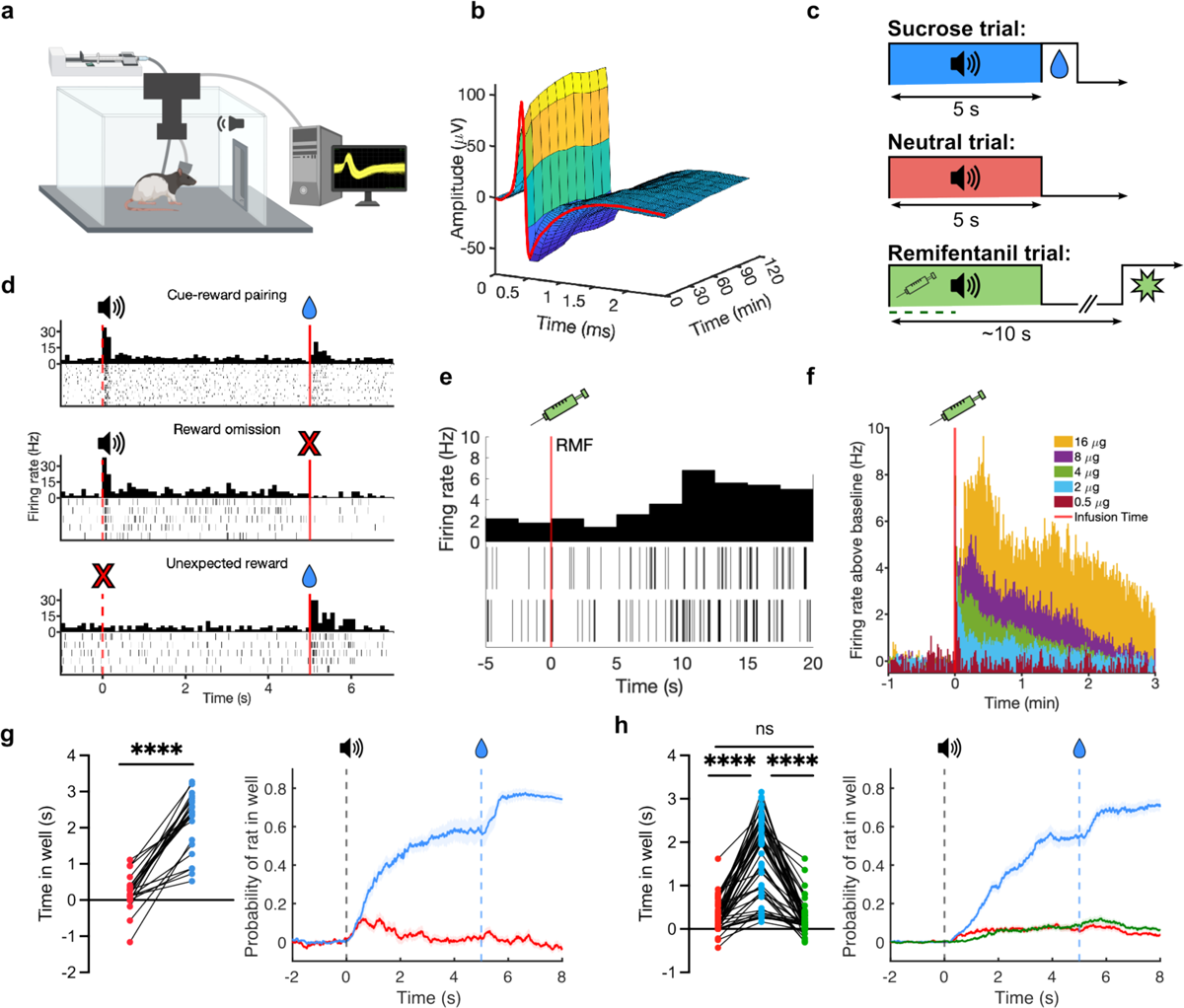
Pavlovian experimental procedure. (**a**) Schematic of operant chamber for training/recording in Experiment 1. (**b**) Example waveform showing stability over a two-hour recording session. Color corresponds to action potential amplitude. (**c**) Pavlovian conditioning of three 5-s tones predicting sucrose (blue), remifentanil (RMF, green), or nothing (neutral, red). (**d**) Example responses of putative dopamine responses to cued sucrose reward (top), reward omission (middle), and uncued reward (bottom). (**e**) Rasters and post-stimulus time histogram (PSTH) showing example dopamine neuron firing after RMF infusion (2 infusions of 4 μg/kg RMF) with onset of RMF effect at ∼10 s post-infusion. (**f**) PSTH for neuronal data of onset and duration of effect of RMF at doses of 0.5 to16 μg/kg/infusion (activation terminates by ∼2.5 min for 4 μg/kg). (**g**) Total time (left) and probability (right) of rat in sucrose reward well during sucrose and neutral cue presentations in opioid-naïve rats (*n* = 22, Wilcoxon test, *z* = −2.105, *p* < 0.0001). Lines and shades represent means ± SEM. (**h**) Same as *h* but in opioid-exposed animals after sucrose, RMF, and neutral cues (*n* = 50, Friedman test, *F* = 75.36, *p* < 0.0001).

Rats were trained with three trial types in each session, each associating a distinct 5-s auditory cue with a unique outcome (Fig. 1c). The dopamine responses to oral sucrose and IV RMF rewards cannot be directly compared, as responses to sucrose reward are precise and discrete (time-locked within the first 1 s following delivery) (Fig. 1d), whereas the firing response to RMF is diffuse, delayed, and variable, peaking and diffusing over tens of seconds (Fig. 1., e and f). However, the model we test predicts that RMF inevitably evokes RPEs that uniquely escalate to maximal drug-cue response, compared to natural-reward cue, regardless of the specific quantity of the RMF used, so such matching is unnecessary. Thus, we simply chose sucrose volumes and RMF doses that are effective reinforcers (Panlilio and Schindler 2000). Cue 1 (henceforth ‘sucrose cue’) was immediately followed by the delivery of a bolus of sucrose at the designated well (40 µL). Cue 2 (‘RMF cue’) was presented simultaneously with activation of the infusion pump for IV RMF delivery (4 µg/kg/infusion). Cue 3 (‘neutral cue’) resulted in no consequence. A subset of rats (n = 4) did not receive any RMF (opioid-naïve comparison group) and were only presented with sucrose and neutral cues. Following RMF infusion, the drug was allowed to clear for approximately 110-320 s (mean = 173 s) based on the observed timecourse of the RMF effect on neuronal firing (Fig. 1f) before initiation of the next trial. Successful discrimination of cue identities was demonstrated by different cue-induced responding, showing significantly greater probability of entering the sucrose well during sucrose vs. RMF or neutral cues (Fig. 1, g and h).

Putative dopamine neurons were identified using hierarchical clustering that has been shown to accurately discriminate genetically identified dopamine neurons (Cohen, Haesler et al. 2012, Eshel, Bukwich et al. 2015, Eshel, Tian et al. 2016, Sadacca, Jones et al. 2016) based on unit activity during sucrose trials (Fig. 2a), and refined by k-means clustering (Supplementary Fig. 1). Identification based on waveform properties and D2 inhibition of select units was more conservative, but otherwise largely agreed with these results (supplementary note 2) (Supplementary Fig. 2).

**Fig. 2.**
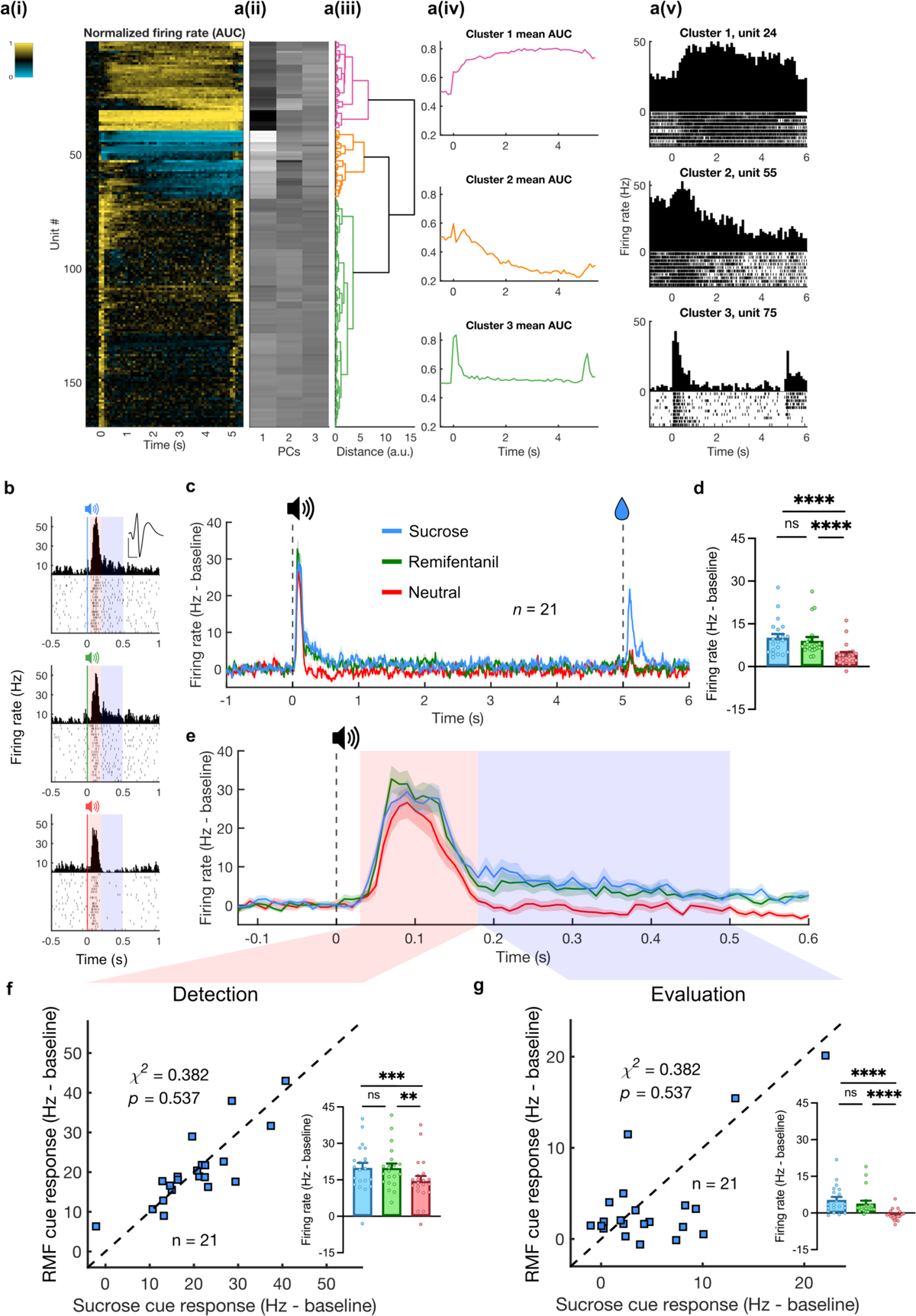
Similar within-neuron dopamine firing responses to sucrose and RMF predictive cues. (**a**) Dopamine neuron identification. (i) Heatmap showing functional activation of each unit (rows) aligned to sucrose cue onset (0 s). Activity normalized using area under receiver-operator curve (auROC) method, compared against baseline. Scales from 0 to 1. (ii) First three components extracted via PCA. (iii) Dendrogram showing results from hierarchical clustering. (iv) Mean auROC for units classified into each cluster. (v) PSTH and raster plots from example units for each cluster. (**b**) Example responses from a single unit to sucrose (blue, top), RMF (green, middle), and neutral (red, bottom) cues. Red and blue shaded regions indicate distinct phases of cue response reflecting detection (30 – 180 ms) and valuation (180 – 500 ms). Inset waveform represents mean of sorted unit. Scale bars: 50 μV and 500 ms. (**c**) PSTH traces showing average dopamine firing during sucrose (blue, 40 μL), RMF (green, 4 μg/kg), and neutral trials (red). (**d**) Mean baseline-subtracted firing rates during cue response to sucrose, RMF, and neutral cues (Friedman test, F = 31.71, *p* < 0.0001). (**e**) Close-up of cue period from Fig. 2c. (**f**) Scatter plot of individual units’ responses to sucrose (abscissa) and RMF (ordinate) cues during the detection phase. Inset: cue responses during the detection phase (F = 16.29, *p* = 0.0003). (**g**) Individual units’ responses during the evaluation phase presented as in Fig. 2f. Inset: cue responses during the evaluation phase (Friedman statistic = 28.67, *p* < 0.0001).

## Within-neuron responses Pavlovian cues

Contrary to the noncompensable RPE hypothesis of cue reactivity, we found no significant difference in the responses of dopamine neurons to cues predicting sucrose (40 μL) vs. RMF (4 μg/kg) cues (n = 21) (Fig. 2, c and d), despite the rats being able to discriminate between these cues (Fig. 1h). Average firing did not differ between sucrose and RMF cues over the 500 ms following cue onset (Fig. 2, b to d). We found that sucrose, RMF, and neutral cues elicited significant responses in the first 30-180 ms after cue onset (‘detection’ component), but only rewarded cues continued to be excited over the next 320 ms (‘evaluation’ component), consistent with a two-component model (Schultz 2016) (Fig. 2, b to d). However, we found no differences between RMF and sucrose cues in either phase (Fig. 2, e to g). In addition, if RMF-associated cues tended to elicit a greater dopamine response than sucrose cues regardless of dose, then we would expect a greater number of neurons to exhibit a higher firing rate in response to RMF than sucrose cues. However, a within-unit comparison of both responses found no significant difference in the numbers of RMF and sucrose-preferring neurons in both the detection (Fig. 2f) and evaluation (Fig. 2g) phases. Both sucrose and RMF reward were associated with increased dopamine response, but over distinct timescales, with RMF response distributed with less moment-to-moment activation over substantially longer duration (Supplementary Fig. 3).

## Sensitized dopamine responses

We compared units obtained from opioid-exposed rats (n = 77) to those from opioid-naïve rats (n = 22) (Fig. 3) and observed that the average baseline firing rate measured before any opioid administration was greater for opioid-exposed than opioid-naïve units (Fig. 3c). Both opioid-exposed and opioid-naïve units showed greater responses to cues predicting 40 μL of sucrose vs. neutral cues, and a significant, positive response to reward delivery simultaneous with sucrose cue offset, but not to neutral cue offset (Fig. 3, e and f). However, opioid-exposed units showed significantly greater responses to sucrose and neutral cues (Fig. 3, g and h) and sucrose delivery (Fig. 3i) than opioid-naïve units. The amount of prior training measured in previous sucrose trials was similar between opioid-naïve and opioid-exposed groups (Fig. 3d), so it is unlikely that any differences in these populations are attributable to differences in duration or experience in training.

**Fig. 3.**
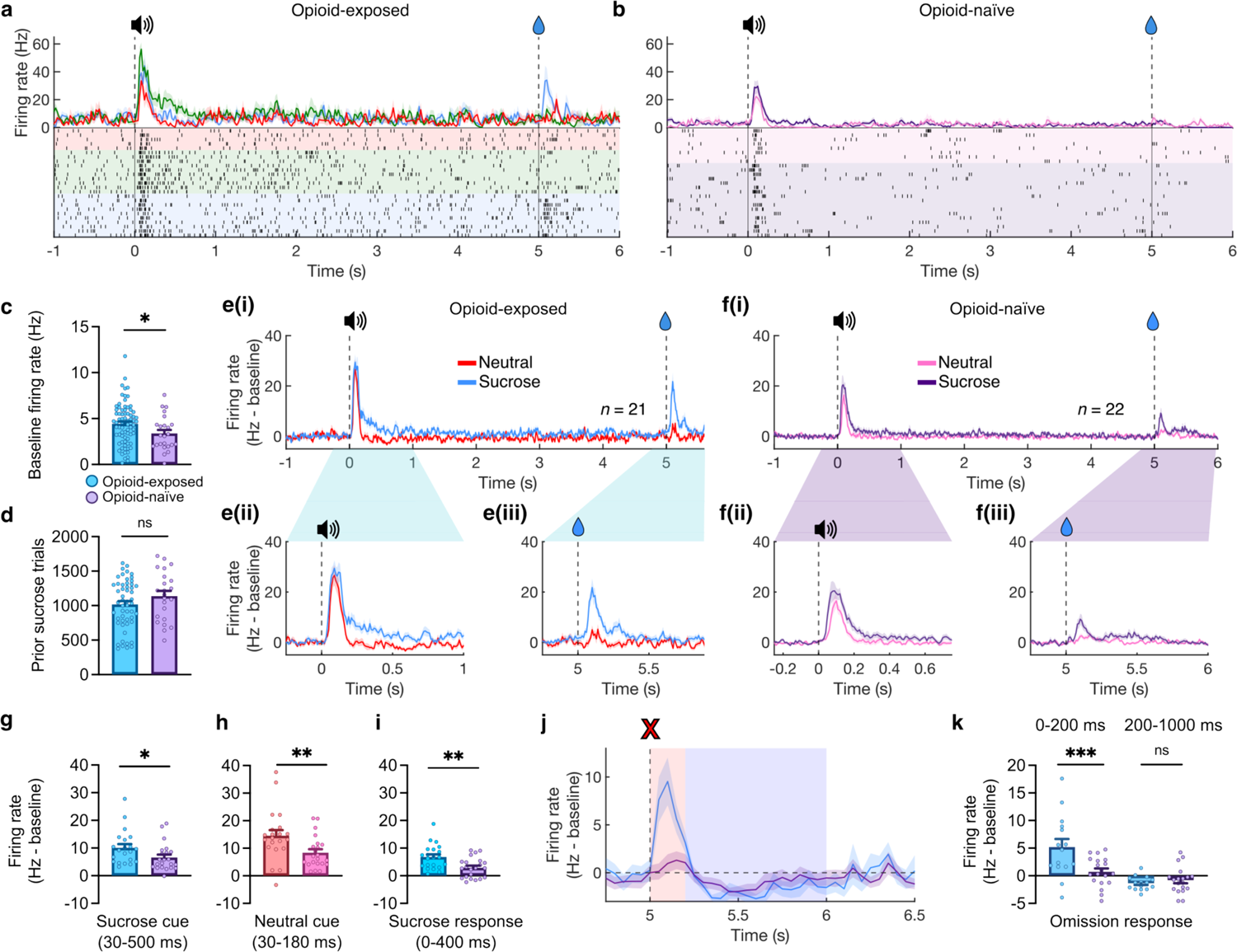
Sensitized dopamine neuron responses in opioid-exposed rats in the Pavlovian procedure. **(a)** PSTH and raster plot examples from a single opioid-exposed unit. Vertical lines indicate cue onset (0 s) and cue offset/sucrose delivery (5 s). Red = neutral trials, green = RMF (4µg/kg), and blue = sucrose (40µL). (**b**) PSTH and raster plot examples from a single opioid-naïve unit. Display similar to *a*, with purple = sucrose and pink = neutral trials. (**c**) Mean firing rates prior to the start of session for each putative opioid-exposed (blue) vs. opioid-naïve (purple) units (*t*-test, *t*(97) = 2.060, *p* = 0.0421). (**d**) Comparison of previous training quantified as number of sucrose trials prior to the recording session (Mann-Whitney test, *n* = 75, *U* = 499, *p* = 0.2586). (**e**) (i) Average dopamine firing during sucrose and neutral (red) trials in opioid-exposed units (same data as Fig. 2). (ii) Close-up of cue response in opioid-exposed units. (iii) Close-up of reward delivery response in opioid-exposed units. (**f**) (i) Average dopamine firing during sucrose and neutral (red) trials in opioid-naïve units. (ii) Close-up of cue response in opioid-naive units. (iii) Close-up of reward delivery response in opioid-naïve units. **(g)**, Comparison of mean cue response to sucrose cues in opioid-exposed (blue) vs. opioid-naïve (purple) units (*n* = 43, *U* = 131, *p* = 0.0145). (**h**) Comparison of mean cue response to neutral cues in opioid-exposed (red) vs. opioid-naïve (pink) units (*n* = 43, *U* = 120, *p* = 0.0064). (**i**) Comparison of mean response to sucrose delivery (0 – 400 ms after cue termination) (*n* = 43, *U* = 118, *p* = 0.0054). (**j**) Trace of dopamine responses at time of reward omission in opioid-exposed (blue) and opioid-naïve units (purple). Red and blue shaded regions reflect positive (0-200 ms) and negative (200-1000 ms) response periods 1 and 2, respectively. (**k**) Comparison of omission firing responses during period 1 and 2 between opioid-exposed and opioid-naïve units (2-way RM ANOVA, Period factor: *F*(1,33) = 25.05, *p* < 0.0001; Exposure factor: *F*(1,33) = 6.087, *p* = 0.0190; Period x Exposure interaction *F*(1,33) = 9.022, *p* = 0.0051).

## Intact natural-reward omission response

In sessions including omission trials (sucrose cue presented without subsequent reward in 10% of trials), both opioid-exposed and opioid-naïve units showed a small negative change in firing at the time of expected sucrose delivery (around 500 ms following cue offset) (Fig. 3j). However, opioid-exposed units exhibited a large positive peak (period 1) before the negative component of the omission response (period 2) (Fig. 3j). Period 1 response likely corresponds to cue offset that also indicates reward availability on rewarded trials (Ferguson, Ahrens et al. 2020, Kalmbach, Winiger et al. 2022), and was significantly greater in opioid-exposed than opioid-naïve units, again reflecting the enhanced salience of the sucrose cue and initial expectation of reward delivery in this group. However, period 2 response was similar between groups (Fig. 3k) suggesting intact negative RPE to natural rewards in opioid-exposed subjects. No negative RPE was observed for RMF omission likely due to the diffuse nature of the RMF-induced firing response.

## Progressive ratio for RMF vs. sucrose

Considering the above results, we sought to verify that our chosen dose of RMF (4µg/kg) was associated with a greater behavioral motivation than sucrose (40 µL) in line with its addictive potential. Therefore, at the end of Experiment 1, rats were trained on a progressive ratio schedule of reinforcement for sucrose (40 µL) and RMF (4 µg/kg). We found a significantly higher breakpoint and total responses recorded for RMF reward vs. sucrose (Supplementary Fig. 4), confirming that the used dose of RMF was of higher motivational value than sucrose.

Results of Experiment 1 show that in opioid-exposed subjects, dopamine neurons exhibit higher baseline activity and generalized enhanced response to both drug and non-drug cues, and to natural rewards. These findings directly challenge the noncompensable RPE model of cue reactivity.

## Operant procedure (Experiment 2)

Experiment 1 is limited by the absence of behavioral measures of drug-cue reactivity, difficulty in directly comparing reward values and corresponding neural responses, and the potential latent pharmacological effect of RMF during subsequent trials. Thus, in a separate group of rats, we conducted Experiment 2 that is based on an operant task whereby distinct discriminative cues predicted the availability of a non-drug or drug reward (trial types A and B, respectively). Cue A was associated with a water reward and Cue B was associated with an identical water reward and a simultaneous RMF infusion (Fig. 4, a and b). As in Experiment 1, a subset of rats remained opioid-naïve, receiving IV saline instead of RMF. This design controls for the sensory and temporal properties of the non-drug reward, and allows direct comparison of the reward values (value(water) < value(water + RMF)). In addition, recording sessions in Experiment 2 occurred in the absence of RMF to avoid any confounding pharmacological effect of the drug. Based on the findings from Experiment 1, we predicted that within-neuron dopamine responses to drug and non-drug rewarded cues will be similar, even in the presence of behavioral drug-cue reactivity.

**Fig. 4.**
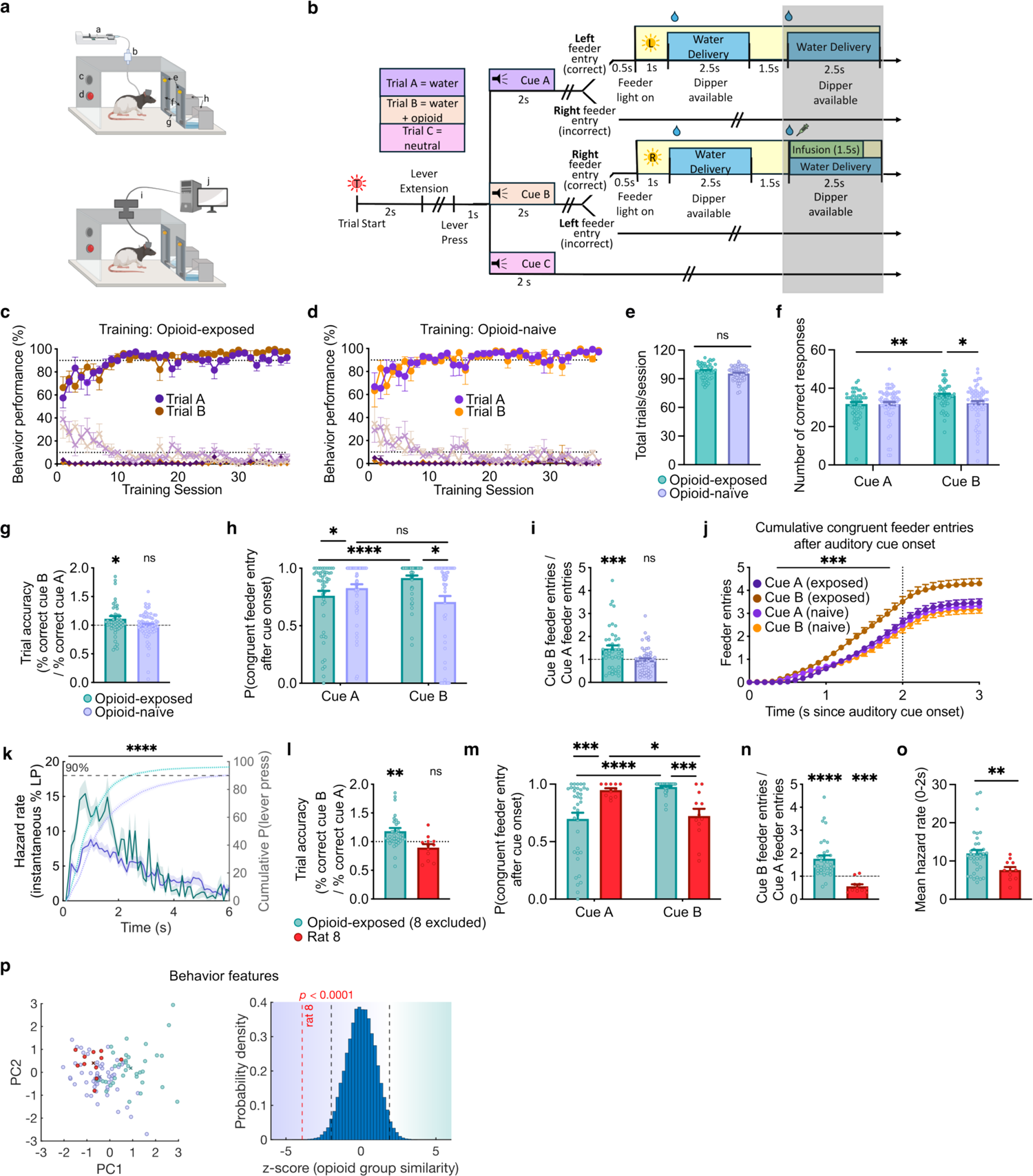
Operant experimental procedure demonstrates behavioral drug-cue reactivity. **(a)** Schematic of experimental setup during training (top) and recording (bottom) sessions in Experiment 2. Boxes equipped with infusion pump (a), swivel (b), speaker (c), trial light (d), feeder lights I, feeders (f), retractable lever (g), dippers (h), commutator (i), and recording computer (j). (**b**) Experimental design. Gray shading indicates omitted reward in block two of recording trials. (**c**) Behavior training results for opioid-exposed animals. Circles indicate % correct responses, Xs indicate % incorrect, and diamonds indicate % not responded to. (**d**) Similar to 4c for opioid-naïve rats. (**e**) Total number of trials completed per recording session for opioid-exposed and opioid-naïve rats (*n* = 106, *U* = 1135, *p* = 0.0975). (**f)** Number of correct responses per session to Cue A and Cue B (2-way RM ANOVA, Cue factor: *F*(1,114) = 4.786, *p* = 0.0307. **(g**) Trial accuracy response index for opioid-exposed and opioid-naïve animals (one-sample *t* tests, Ho: *μ* = 1. Opioid-exposed: *t* = 2.168, *df* = 47, *p* = 0.0353; Opioid-naïve: *t* = 0.8954, *df* = 63, *p* = 0.3740). (**h**) Probability that first feeder entry after cue onset is congruent with cue identity (2-way RM ANOVA, opioid exposure factor: *F*(1,101) = 5.032, *p* = 0.027; interaction *F*(1,101) = 8.621, *p* = 0.0041). (**i**) Ratio of total congruent feeder entries for Cue B/Cue A in both groups (one-sample *t* tests. Opioid-exposed: *t* = 3.343, *df* = 46, *p* = 0.0017; Opioid-naïve: *t* = 0.6371, *df* = 54, *p* = 0.5268). (**j**) Cumulative congruent feeder entries following auditory cue onset (F-test for difference of slopes, *F*(3,156) = 5.482, *p* = 0.0013). (**k**) Left axis: Session-mean of lever press hazard rate. 2-way Mixed effects model, Opioid-exposure factor: *F*(1,108) = 24.04, *p* < 0.0001). Right axis: cumulative % of sessions with lever press at time *t* is shifted to the left in opioid-exposed rats. (**l-n**) As in h-j respectively, but for rat 8 (red) and all other opioid-exposed animals: m (Opioid-exposed: *t* = 3.284, *df* = 36, *p* = 0.0023; rat 8: *t* = 1.730, *df* = 11, *p* = 0.1115); n (interaction *F*(1,47) = 22.4, *p* = <0.0001); o (Opioid-exposed: *t* = 5.352, *df* = 34, *p* < 0.0001; rat 8: *t* = 5.000, *df* = 10, *p* = 0.0005). (**o**) Mean hazard rate for lever press during 0-2 s after lever extension in rat 8 vs. other opioid-exposed rats (*n* = 49, *U* = 91, *p* = 0.0038). (**p**) Left: First two PCs extracted from composite behavior measures for opioid-naïve rats (purple), rat 8 (red filled), and all other opioid-exposed rats (teal). Crosses represent corresponding centroids. Right: Mean difference of rat 8 centroid under true labels (red dashed line) compared to bootstrapped distribution under shuffled labeling shows greater similarity of rat 8 to the opioid-naïve group. Black dashed lines indicate two-tailed 95% threshold of null distribution.

Rats in the opoid-exposed and opioid-naïve groups were required to achieve >90% accuracy on both trial types A and B before proceeding to recording sessions (Fig. 4, c and d). These occurred in a distinct operant chamber where no IV line was attached but a high level of accuracy was generally maintained for both trial types in both groups (Fig. 4f and Supplementary Fig. 5). Our data show enhanced reactivity to the drug cue in opioid-exposed rats as suggested by multiple lines of evidence. In opioid-exposed animals – but not opioid-naïve – the relative trial accuracy index (% correct Cue B responses/% correct Cue A responses) across sessions was biased in favor of Cue B (Fig. 4g). Only port entry following cue termination determined correct vs. incorrect trial classification, so we examined port entries, in the correct trials, during auditory cue presentation. In opioid-exposed rats, the first feeder entry following Cue B onset was more likely to be congruent with the correct feeder entry compared to the first feeder entry after Cue A in the same group and compared to the first feeder entry after Cue B in the opioid-naïve group (Fig. 4h). We recorded the cumulative number of congruent port entries for both cue types in both groups and observed that accumulation of correct entries proceeded at a substantially faster rate for Cue B in opioid-exposed than in all three other conditions, detectable as both greater cumulative entries up to one sec following cue termination (Fig. 4i) and greater slope during cue presentation (Fig. 4j). Behavioral reactivity was also seen in the more expedient lever-pressing response following lever extension in the opioid-exposed group. Instantaneous probability of a lever press (only one lever press was observed per trial, as the lever was immediately retracted) after lever extension, was quantified as hazard rate (Hamid, Pettibone et al. 2016). This was significantly higher in opioid-exposed vs. opioid-naïve rats over the first two seconds following lever extension (Fig. 4k), resulting in a shorter time to 90% cumulative probability of lever press in the opioid-exposed (∼2 s) vs. opioid-naïve group (∼6 s). These results demonstrate both selective drug-cue reactivity as well as generalized increase in motivation in the drug-exposed group.

While we observed behavioral drug-cue reactivity overall for the opioid-exposed group, one rat (rat 8) was a notable exception to this trend. This rat did not display drug-cue reactivity and showed more preference to the non-drug cue across all behavioral measures (Fig. 4, l to o), even though it completed a similar number of trials and earned similar number of rewards to other opioid-exposed and opioid-naïve rats (Supplementary Fig. 5, e and f). The behavioral profile of rat 8 was more like the opioid-naïve than the rest of the opioid-exposed rats (Fig. 4p). Thus rat 8 could serve as a case-control of opioid exposure for the role of dopamine firing in drug-cue reactivity.

## Dopamine firing responses to operant cues

With behavioral evidence of drug-cue reactivity, we then examined whether this behavior would be reflected in the activity of midbrain dopamine neurons. Dopamine neuron identification (n = 69) was like Experiment 1 (Supplementary Fig. 6). We performed our analyses both excluding units recorded from rat 8 due to the absence of behavioral cue reactivity (Fig. 5), and including the entire dataset (Supplementary Fig. 7).

**Fig. 5.**
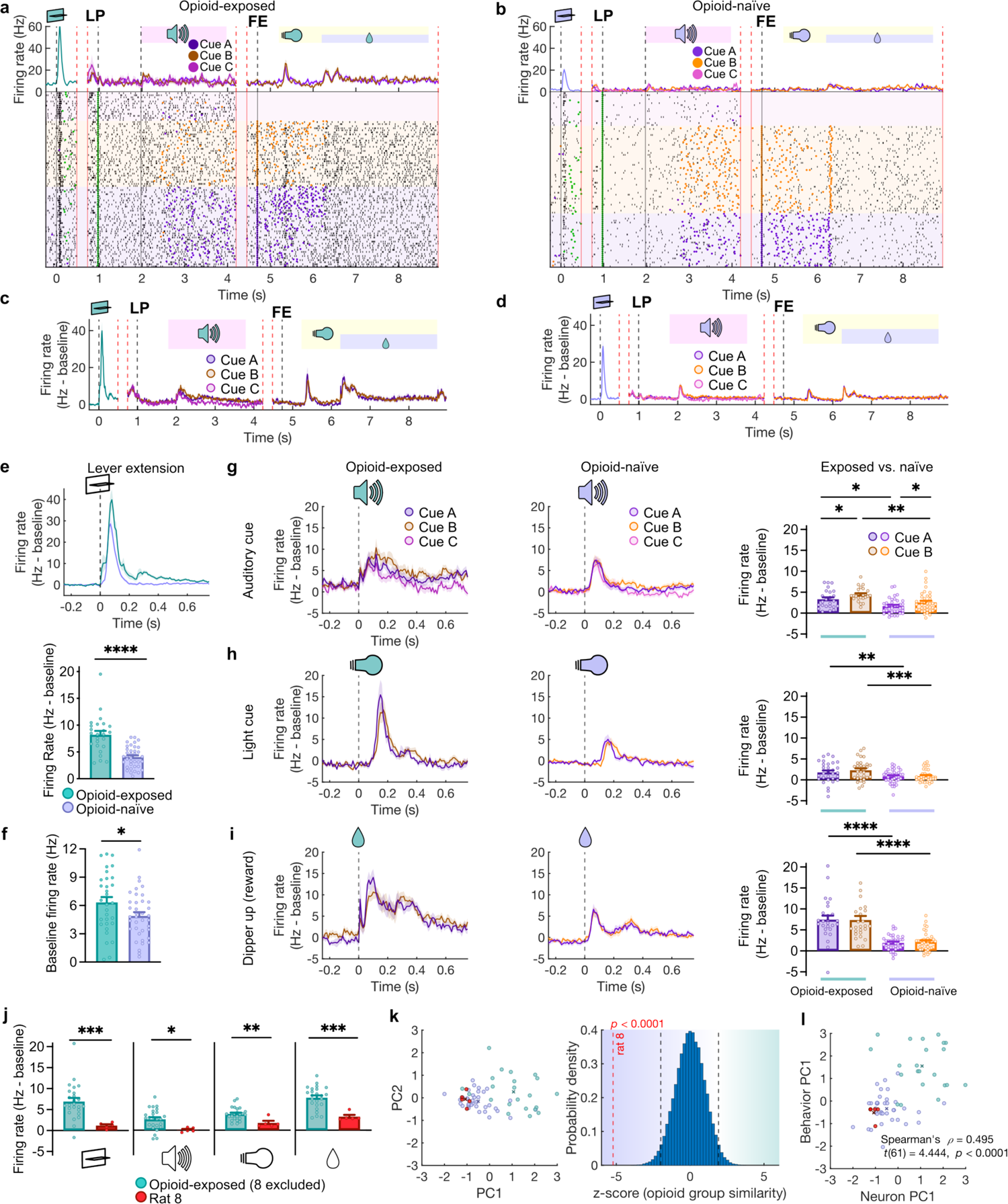
Sensitized dopamine responses to drug and non-drug cues in opioid-exposed rats with behavioral drug-cue reactivity. (**a**) PSTH and raster plot examples from a single opioid-exposed unit. Vertical red lines indicate timeline discontinuity due to variable intervals within trials. Vertical black lines indicated discrete events: lever extension, lever press, and rewarded feeder entry. Purple trace includes all trials, which subsequently split based on trial identity to corresponding colors. Purple and orange dots indicate left and right feeder entries respectively. Green dots indicate lever presses. (**b**) Similar to a but for an opioid-naive unit. (**c**) Mean PSTH from opioid-exposed units displayed like in a. (**d**) Mean PSTH from opioid-naïve units. (**e**) Top: PSTH of baseline-subtracted firing rate around lever extension for opioid-exposed and opioid-naïve units. Bottom: mean firing rate for both groups (Mann-Whitney test, *n* = 62, *U* = 124, *p* < 0.0001). (**f**) Baseline firing rate for opioid-exposed (teal) and opioid-naïve (purple) units (t-test, *df* = 67, *t* = 2.117, *p* = 0.0379). (**g**) Responses of opioid-exposed and opioid-naïve units to the auditory cues. Right: mean firing rates (2-way RM mixed-effects analysis, Cue factor: *F*(1,64) = 10.79, *p* = 0.0017; Exposure factor: *F*(1,65) = 8.157, *p* = 0.0058). (**h**) Responses of opioid-exposed and opioid-naïve units to light cue. Right: mean firing rates (trial-type factor: *F*(1,60) = 2.110, *p* = 0.1515; Exposure factor: *F*(1,62) = 11.96, *p* = 0.0010). (**i**) Responses of opioid-exposed and opioid-naïve neurons to water reward delivery. Right: mean firing rates (trial-type factor: *F*(1,61) = 0.1357, *p* = 0.7139; Exposure factor: *F*(1,62) = 38.41, *p* < 0.0001). (**j**) Mean responses of neurons from rat 8 (red) vs. all other opioid-exposed units in response to lever extension (*t*(27) *=* 3.695, *p* = 0.0010), auditory cue (A and B; *t*(27) *=* 2.915, *p* = 0.0071), light cue (A and B; *t*(27) *=* 2.609, *p* = 0.0355) and reward delivery (A and B; *U* = 8, *p* = 0.0008). (**k**) Left: First two PCs extracted from composite firing measures for opioid-naïve rats (purple), rat 8 (red filled), and all other opioid-exposed rats (teal). Crosses represent corresponding centroids. Right: Mean difference of rat 8 centroid under true labels compared to bootstrapped distribution under shuffled labeling plotted like in Fig. 4p shows greater similarity of rat 8 to the opioid-naïve group. (**l**) Scatter plot of PC1 from neuron firing activity analysis in K vs. PC1 from behavior analysis from Fig. 4p shows moderate positive correlation between behavioral cue reactivity and and dopamine firing response.

Lever extension resulted in the largest dopaminergic firing response compared to other trial events, and was significantly higher in the opioid-exposed (n = 31) compared to opioid-naïve rats (n = 38) (Fig. 5, a to e). The discriminative auditory cues provoked significant phasic dopamine responses across all trial types, including a larger response to Cue B than Cue A, in both opioid-exposed and opioid-naïve groups (supplementary note 3) (Fig. 5g). However, the firing responses to both Cue A and Cue B were larger in the opioid-exposed than opioid-naïve rats (Fig. 5g). After termination of the auditory cue and correct feeder entry, the light cue that confirmed correct responding and signaled upcoming reward delivery also resulted in significant dopamine activity. In opioid-exposed rats, this light-cue response was significantly greater than in the opioid-naïve comparison group and there was no within-neuron difference in the mean response between the light cues predicting water vs. water + RMF/saline within each group (Fig. 5h). Overall, these results show that dopamine firing responses to cues associated with a (water + drug) reward were similar to water-only reward in the opioid-exposed group, and both were higher than opioid-naïve group.

Finally, when the water reward was delivered, the phasic dopamine response was also greater for neurons in opioid-exposed than opioid-naïve rats (Fig. 5i). There was no such difference between the trial types A and B in either opioid-exposed or opioid-naïve rats (Fig. 5i). No negative RPEs were observed for the omitted second water reward in either opioid-naïve or exposed groups (Supplementary Fig. 5, h and i), likely due to its predictability. Of note, baseline firing measured during ITIs was again significantly higher in dopamine neurons from opioid-exposed rats, like in experiment 1 (Fig. 5f).

## Relationship of firing to cue reactivity

We compared the activity of dopamine neurons from rat 8, which did not show cue reactivity despite an extensive history of RMF self-administration (achieved >90% accuracy during training sessions), to the rest of the opioid-exposed population. Rat 8’s neurons consistently showed significantly lower firing across all trial events, and its overall neural response profile was more like the opioid-naïve than the rest of the opioid-exposed units (Fig. 5, j and k). These results are consistent with a role of enhanced phasic dopamine responses in cue reactivity. In support of the relationship between dopamine responses and behavioral drug-cue reactivity, we found a significant moderate, positive correlation between the first PCs of neuronal firing and behavioral cue reactivity from the sessions in which they were recorded (supplementary note 4) (Fig. 5l). The difference in dopamine firing between the opioid-exposed and naïve groups or rat 8 was not reflected in the number of trials completed or rewards received (Supplementary Fig. 5, e and f).

## Discussion

This study challenges the prevailing hypothesis of cue reactivity that opioids act as noncompensable positive RPEs, driving selective enhancement of dopamine response to drug-related cues. We found a non-selective effect of opioid exposure on dopamine neuron activity, particularly elevated baseline firing rate, and enhanced phasic responding across cue and reward types and in multiple distinct experimental setups. There was largely no difference in the magnitude of dopamine firing responses to cues predicting an opioid drug compared to a natural reward, even when the drug reward was clearly of higher value (drug + water vs. water). These findings contradict the widely held RPE-based assumptions of abnormal cue reactivity in the addiction literature, which suggest that dopamine neurons systematically encode lower values for natural reward cues than drug cues, or even natural reward cues in non-drug-exposed individuals (Redish 2004, Kalhan, Redish et al. 2021).

An alternative to the noncompensable RPE hypothesis of cue reactivity that our findings support is that exposure to opioids, rather than accumulating value on drug-specific cues (cached value) and thus selectively enhancing dopaminergic responses to these cues, results in increased excitability of midbrain dopamine neurons, which manifests as a generalized reinforcement gain. This gain causes enhanced but reward-dependent dopamine response to all salient cues that occur in the same spatiotemporal context as drug use including non-drug predicting and neutral cues (supplementary note 5). Such non-specific cue reactivity has been shown clinically in smokers, where reactivity to drug and non-drug cues covary significantly (Mahler and de Wit 2010). One possibility for such an effect is that drug-induced dopamine release is sustained over minutes, unlike the physiologic brief and discrete dopamine release in response to natural rewards, and hence causes long-lasting changes in signaling cascades and gene expression that outlast the time course of dopamine release. These changes, which normally reinforce drug-related cues and behaviors, overlap with and thus also reinforce other spatiotemporally overlapping cue-reward associations in the animal’s environment, effectively causing a generalized increase in the gain on future reward expectation in the RPE calculation. Prior literature supports the finding of increased dopaminergic response to drug cues (Diana, Muntoni et al. 1999, Zijlstra, Booij et al. 2008, Lefevre, Pisansky et al. 2020, Samaha, Khoo et al. 2021, Leyton 2022) but see (Volkow, Wang et al. 2011, Trifilieff and Martinez 2014) (supplementary note 6). Further, opioid use is linked to increased excitatory and reduced inhibitory synaptic transmission onto midbrain dopamine neurons (Saal, Dong et al. 2003, Moussawi, Ortiz et al. 2020), which could contribute to increased dopamine neuronal excitability. The dopamine-dependent psychomotor sensitization after opioids (Steketee and Kalivas 2011) and increased dopamine neuronal firing to acute opioids after chronic opioid exposure (Diana, Muntoni et al. 1999) also point towards increased excitability of dopamine neurons. This is also supported by the observed increase in baseline firing rate in the opioid group. Tonic firing has been theorized to reflect real-time value estimates (Niv, Daw et al. 2007, Mohebi, Pettibone et al. 2019), which concords with a general motivational sensitization reflected in the increased hazard rate of lever pressing. In addition, the generalized reinforcement gain model is more consistent than the noncompensable RPE model with prior studies that failed to show overwhelming drug preference in animal models (supplementary note 7) (Ahmed, Lenoir et al. 2013, Caprioli, Zeric et al. 2015, Chow and Beckmann 2021). Further, this alternative model of non-selective reinforcement gain through sensitization of dopamine function is consistent with prior literature showing that activation of dopamine circuitry by drugs or medications heightens sensitivity to rewards that are not directly associated with the pharmacological agent. For example, cue-induced responding for natural rewards can be increased by exposure to opioids, amphetamines, or cocaine (Robbins 1978, Taylor and Robbins 1984, Wyvell and Berridge 2001, Ostlund, LeBlanc et al. 2014) including in a drug-free state and in a distinct context (Wyvell and Berridge 2001). Similarly, patients taking dopamine-augmenting medications for movement disorders can develop addiction-like compulsive behavioral problems as a side effect (Voon, Hassan et al. 2006, O’Sullivan, Evans et al. 2009).

Our results show a disconnect, in the opioid-exposed group, between the cached value of reward cues measured with dopamine firing and their actual value reflected in the drug-cue reactivity. However, our data also show a strong relationship between enhanced dopamine firing and drug-cue reactivity. Thus, although enhanced excitability of dopamine neurons may have a key role in promoting reactivity, behavioral selectivity is not generated by a difference in dopamine firing to drug vs. non-drug cues. Compelling cross-species evidence suggests that mammalian reward learning balances elements of model-free (e.g., temporal-difference) and model-based learning (Daw, Gershman et al. 2011, Doll, Simon et al. 2012). It is possible that the non-specific dopamine enhancement we observe here could promote behavioral activation which is sculpted to promote drug-seeking by causal information supplied by structures implicated in model-based learning, such as the orbitofrontal cortex. A recent report has suggested a similar role for the orbitofrontal cortex in promoting drug addiction by interfering with natural reward consumption (Tan, Browne et al. 2024). Alternatively, it is also possible that behavioral selectivity is dictated by regulation of dopamine release distally at axon terminals, which may be regulated independently from somatic firing (Mohebi, Collins et al. 2023).

However, other reports have shown that closer matching of somatic and axonal dopamine dynamics substantially alleviates the discrepancy (Azcorra, Gaertner et al. 2023). In our data, drug and non-drug cues were learned in a single context, so it is unclear whether the enhanced dopamine response to non-drug cues and rewards is spatiotemporally-specific.

Because we hypothesize that the observed enhancement is a result of aberrant learning around drug use, we further postulate that our findings are drug-context specific and thus carry important implications for the clinical setting and explain the high relapse rate of patients discharged to their home environment where they experienced drug use. Non-drug cues learned in proximity to drug acquire similar cached value to drug cues and remain a clinically important challenge as increased dopamine firing and release are critical for relapse (Kalivas and Volkow 2005). In summary, results from this study warrant revised models of dopaminergic function in drug addiction and provide critical insights into the neurobiology of cue reactivity with potential implications for relapse prevention.

## Methods

### Subjects

All experimental procedures were conducted in accordance with the guidelines of the National Institutes of Health Guide for the Care and use of Laboratory Animals and approved by the Animal Care and Use Committee of the University of Pittsburgh (Experiment 1) and the National Institute on Drug Abuse (Experiment 2). Experiment 1 involved 11 adult Long-Evans rats (age 9-15 weeks, mean = 10.5 weeks, 9 males, 2 females) with 7 rats in the opioid-exposed group and 4 rats in the opioid-naïve group. Experiment 2 involved 11 male Long-Evans rats (age 9-12 weeks) with 5 rats in the opioid-exposed group and 6 rats in the opioid-naïve group.

### Electrode surgeries

For Experiment 1, rats were implanted with 32-channel drivable electrode bundles targeting left VTA (ML = −0.50 mm, AP = −5.40 mm, DV = −7.40 mm), protected by plastic caps.

Electrodes were constructed from 25µm formvar-insulated NiChrome wires (A-M Systems Carlsborg, WA) as previously described (Takahashi, Batchelor et al. 2017). The wires were bundled in two 27-gauge cannulas centered 740 um apart and mounted in a custom 3D-printed microdrive. Prior to implantation, wires were trimmed to 1-2 mm, spread to allow ≥ 25 µm between them, and gold-plated to an impedance of 400-700 kΩ (at 100 Hz). Surgeries were performed under isoflurane with aseptic technique, and 0.5 mg/kg Carprofen was provided for two days for pain management. For two weeks, oral cephalexin and topical Neosporin were also provided.

For Experiment 2, rats were implanted with similar electrodes targeting the VTA. In this experiment, most rats had 8-channel electrodes, while one had 32 channels.

### Catheter surgeries

For Experiment 1, after two weeks of recovery from electrode surgery, indwelling intravenous catheters were implanted in the right jugular vein and tunneled to a vascular access port on the rat’s back (Instech) as previously described (Moussawi, Zhou et al. 2011), followed by a third week for recovery, in which the catheter was flushed daily with 0.9% saline and gentamicin (4.25 mg/kg). Thereafter, catheters were flushed daily with a solution of saline, heparin and enrofloxacin, and tested weekly with propofol to ensure patency.

For Experiment 2, IV catheter implantation was done as in Experiment 1. Both cohorts underwent similar long-term treatment with saline and antibiotics, and regular propofol testing to ensure maintained patency.

### Pavlovian procedure in Experiment 1

Rats were water restricted (30-60 min access following behavior each day) and trained in a customized operant chamber equipped with an overhead tether for IV fluid delivery, a well connected to solenoid valves for delivery of oral sucrose (10% w/v) and vacuum for removal, and a speaker opposite the port to play auditory cues. During training, the beginnings and ends of sessions were marked by a house light turning on and off respectively. Remifentanil was prepared each day from stock stored at −20 °F.

Training consisted of three trial types with distinct 5s auditory cues. In sucrose trials, a 5 s 8 kHz tone was immediately followed by sucrose dispensed at the well with a vacuum activated later to remove any unconsumed liquid between trials. In remifentanil trials, a noiseless IV pump dispensing RMF was activated simultaneously with onset of a 5 s 12 kHz tone to minimize the delay between cue onset and onset of pharmacological effect of RMF. In neutral trials, a 5 s cue rapidly cycling between 2.5, 3, 3.5, and 4 kHz was followed by no consequence. ITIs between trials were selected pseudorandomly and lasted 11-150 s, (mean 68.7 s or 69.9 s, following sucrose and neutral trials) or 110-320 s (mean 173 s, following RMF trials). Sucrose and RMF rewards (sucrose from solenoid and remifentanil IV pump) were omitted in a randomly selected 10% of trials. Training sessions consisted of one or two blocks per session with 25-50 trials per training block (typically 2:2:1 ratio of RMF, sucrose, and neutral trials) with occasional sucrose or RMF reward omission (∼10% of trials). Opioid-naïve rats were trained only on sucrose and neutral cues.

### Progressive ratio testing

Six male Long-Evans rats trained in the Pavlovian procedure were subsequently trained on a progressive ratio task. Training began with an FR1 schedule for 4 <u>μ</u>g/kg IV RMF reward paired with the familiar drug cue per lever press. Each session lasted one hour, and training continued for at least five sessions, and until each rat received at least 50 infusions in one session.

Progressive ratio testing was carried out with exponentially escalating response requirements (Richardson and Roberts 1996) (steps = 1, 2, 4, 6, 9, 12, 15, 20, 25, 32, 40, 50, 62, 77, 95, 118, 145, 178, 219, 268, 328, 402, 492, 603, and 737). Each session was terminated after three hours or a period of 30 minutes with no reward. Recorded scores are the highest of ∼3 PR sessions per animal. After RMF testing, rats were trained to press a lever located on the other side of the operant chamber for 40 μL oral sucrose. Training began in FR1, with each lever press resulting in a familiar sucrose cue, and delivery of liquid at the same location as in Pavlovian conditioning. Once again, training was maintained for at least five sessions, or until at least fifty rewards were collected in a one-hour session. Subsequently, progressive ratio testing for sucrose reward was carried out according to the same schedule, and recorded scores represent the highest of ∼3 sessions per animal.

### Operant procedure in Experiment 2

In Experiment 2, rats were water restricted and trained with trials resulting in natural reward, (natural+drug) reward, and no reward. Natural reward consisted of water, and the drug reward (4 μg/kg IV RMF) was paired with an identical water reward. All operant trials began identically with illumination of an overhead trial light, followed two seconds later by extension of the lever. A lever press resulted in lever retraction and, one second later, by one of three discriminatory melodic auditory cues played to indicate the trial type. Note that due to the proximity of lever presses to lever extension, firing responses during lever press are difficult to compare, as neural responses do not return to baseline before the behavioral event (Supplementary Fig. 5d). The auditory cues were pseudorandomized and signaled the location and identity of the reward (Cue A, “siren”: water only, left port; Cue B “white noise”: water + RMF/saline, right port; Cue C “3 kHz beeping 2x/s”: no reward - neutral trial). A port entry during cue presentation was inconsequential irrespective of congruency with correct feeder side. A correct port entry after the offset of Cue A resulted 0.5 s later in the illumination of a light cue within the feeder port for the whole duration of reward delivery. One second after the light cue was turned on, two water rewards were delivered via a motorized dipper (Coulbourn H14-05); each water reward was 40 µl and was delivered over 2.5 s with 1.5 s interval between the rewards during which the dipper was retracted outside the feeder port (Fig. 4, a and b). A correct port entry after the offset of Cue B resulted in a similar cascade of cues and rewards as to Cue A response (including 2 x 40µl water rewards) but with the additional activation of an infusion pump to deliver IV RMF (4µg/Kg/50µl infusion) or saline at the onset of the second water reward delivery (Fig. 4b). The intertrial interval (ITI) for correct trials A and B types was 60 ± 15 s. Incorrect response to Cue A or B resulted in a prolonged ITI (120 ± 15 s). If rats failed to respond to Cue A or B within 10 s, a new trial was initiated and the lever re-extended. Type C cues were inconsequential, and Cue C offset was followed by a shortened ITI (30 ± 10 s). Each training session was 4 hrs. Type A, B, and C cue trials were pseudorandomized from a list with typically 1:1:1 ratio. Refer to Fig 4 a,b for detailed schematic of recording chamber arrangement.

### Single unit recordings and spike sorting

During recording, electrodes were connected to a headstage tether which was prevented from tangling with the IV tether by a custom motorized commutator in Experiment 1 (Plexon, inc.). Single unit recordings were acquired with the Plexon OmniPlex system. Signals were digitized at 40 kHz and band-pass filtered at 0.1-7500 Hz, then digitally high pass filtered at 0.77 Hz. Spike data was separated from LFPs by an additional 150 Hz high-pass filter. Between sessions, electrodes were advanced 40-80 um. Spikes were isolated and single units sorted manually in Offline Sorter (Plexon, Inc.) with an SNR cutoff of 3:1 (*σ_spikes_*^2^/*σ_noise_*^2^).

### Recording sessions in the Pavlovian procedure (Experiment 1)

Single-unit recordings in Experiment 1 occurred throughout the behavioral training sessions described above.

### Recording sessions in the operant procedure (Experiment 2)

Single-unit recordings in Experiment 2 occurred in distinct sessions from the training sessions. Recording sessions were generally similar to the training sessions with the following exceptions: The recording sessions involved 2 blocks during which the IV infusion was omitted for both groups, and recordings occurred in a customized operant chamber in a different location than training boxes where no IV line was attached. Block one was identical to training except the lack of IV infusions and lasted until the rats correctly responded to twelve rewarded trials. Once this requirement was met, rats advanced to block two, which was identical to block one except for the omission of the second reward in rewarded trials. This design allows for examination of negative RPEs in opioid-exposed and opioid-naïve groups. A recording session usually consisted of a maximum 72 total correct type A and B trials, or 90 min duration, with typically 2:2:1 ratio of types A, B, and C trials). The ITIs were shorter than training sessions (correct response to Cue A/B: 35 ± 5 s; incorrect response to Cue A/B: 50 ± 10 s; Cue C: 20 ± 5 s). Between recording sessions, rats were retrained for at least 2 sessions to prevent behavioral extinction and were required to achieve >90% accuracy on both trial types A and B before a new recording session was carried out.

### Dopamine unit identification

Waveform properties were calculated from the average waveform extracted from each unit. The amplitude ratio was defined as the ratio of the difference to the sum of the first positive peak detected before the first negative peak and the first negative peak. If the first peak was negative, (no preceding positive peak), the first positive peak was taken to have amplitude of 0. The half duration was defined as the time from the first negative peak to the next positive peak, or the end of the waveform (at 1.1 ms) if no such positive peak was detected (Takahashi, Batchelor et al. 2017, Takahashi, Stalnaker et al. 2023). After screening for cue responsiveness in Experiment 1, 225 units were considered for further analysis. To identify putative dopamine neurons, we used an activity-based clustering approach that identifies dopamine neurons based on canonical phasic responses to conditioned cues and reward, which has been shown to accurately discriminate genetically identified dopamine cells (Cohen, Haesler et al. 2012, Eshel, Bukwich et al. 2015, Eshel, Tian et al. 2016).

Each of the units’ spikes was isolated in a period 0.5s before and 5.5s after each sucrose cue, and a normalized response was calculated with the auROC method. The first three principal components of these normalized responses were clustered hierarchically into three groups using Ward’s distance metric (cluster 1 “excited”, n = 39, cluster 2 “inhibited”, n = 30, cluster 3 “phasic”, n = 165). The group that displayed classic phasic “RPE-like” dopamine activity and was labeled as putative dopamine. Within the putative dopamine cluster, we observed a subpopulation with delayed peak firing, highly labile firing, and sustained inhibition to reward. We calculated peak firing time as the time of highest activity 50-1000 ms after sucrose cue onset, binned at 20 ms, firing variability as the coefficient of variation of ISIs recorded throughout the sessions excluding periods 0-500 ms after cue/reward events (where phasic activity is expected), and inhibition as (R-B)/(R+B) where R was the mean response 1000-2000 s after reward delivery and B was the session baseline firing rate. We performed PCA on normalized log(peak firing time), log(ISI CV), and inhibition and used K-means clustering on the first two principal components (*k* = 2, Euclidean distance, 20 replicates) to separate this group (n = 21) from canonical dopamine neurons. We excluded any units exhibiting baseline firing > 12 Hz from this clustering (n = 39). Finally, 6 units were removed manually (Supplementary Fig. 9 a) leaving 99 putative dopamine neurons in our analysis. Since dopamine units were not directly genetically identified, we cannot rule out the possibility that some excluded units are dopamine units that deviate from canonical dopamine activity. However, if such neurons are present, they are not well-characterized in prior literature, and their activity is not predicted by conventional models of dopamine activity. Seventy-five of the putative dopamine neurons were recorded from rats with prior opioid exposure (opioid-exposed), while the remaining 24 were recorded from comparison rats which received no opioids (opioid-naïve). There was no significant difference in the waveform properties between the opioid-exposed and opioid-naïve groups (Supplementary Fig. 8a), and putative dopamine neurons exhibited classic phasic response to cues, with firing response to cues peaking at ∼100 ms (Fig. 2 and Supplementary Fig. 8b) in both groups. Further, there was no difference in the bursting behavior in terms of frequency of bursts, percentage of spikes in bursts, or distribution of burst sizes (Supplementary Fig. 8, c to e).

Identification in Experiment 2 was achieved with a similar approach. Since we lacked a simple cue-reward pairing, a composite cue-reward period was created by appending activity recorded during around initial lever extension (−250 to 1000 ms) to activity around reward delivery (−250 to 500 ms). Hierarchical clustering on auROC signals again yielded three groups exhibiting sustained excitation (n = 55), inhibition (n = 39), and phasic firing (n = 187). After eliminating those units in the ‘phasic’ cluster with high baseline firing rate (> 12 Hz; n = 75), we again observed a subset of units with sustained inhibition. Peak firing time, ISI CV, and inhibition were calculated as in Experiment 1, except that the inhibition response was the lower of mean firing 0-1000 ms before or after reward delivery. PCA as in Experiment 1, followed by K-means clustering (*k* = 2, Euclidean distance, 20 replicates), eliminating 30 units. A further three units were eliminated manually (Supplementary Fig. 9 b) and the remaining units were classified as putative dopamine (n = 69).

### Rat 8 similarity analysis

To test the similarity of rat 8 behavior to the opioid-naïve vs. opioid-exposed rats, we performed a PCA on four behavioral measures (cumulative feeder entry ratio, correct trial index, mean hazard rate, and congruent feeder entry ratio) and retained the first two PCs. We randomly sampled with replacement from the naïve sessions and the opioid sessions (excluding those from rat 8) and computed the difference in Euclidean distances between these points and the centroid of rat 8’s behavioral data (mean distance between opioid-naïve sessions to centroid of rat 8 from mean distance between opioid-exposed sessions and centroid of rat 8). Averaging the differences in distances obtained by 2000 such samples allowed us to generate a bootstrapped estimate of the mean similarity to the naïve and exposed groups with negative values reflecting greater similarity of rat 8 behavior to the naïve group, while positive values reflecting greater similarity to the exposed group. We generated a null distribution by randomly relabeling the behavioral data from both groups (excluding rat 8) with the same proportion of exposed and naïve sessions and computing 50,000 relabeled similarity scores (Fig. 4p). Statistical significance was assessed by computing the percentile of the true similarity score based on the null distribution.

To test for similarity of single unit activity, we performed PCA on four measures of spiking activity (mean response to lever extension, auditory cues A and B, light cues A and B, and rewards A and B). Subsequent analysis on the first two PCs was performed identically to behavior data.

### Data processing

Peristimulus temporal histograms were constructed from raw spike trains with bins of 10 ms width, except where otherwise noted. Smoothing was carried out by convolving the binned data with a causal exponential kernel over 200 ms (Thompson et al, 1996). Lines and surrounding shaded areas in traces show mean ± SEM of data.

In Experiment 1, well entries and exits were recorded and used to calculate the PSTH of probability of presence in the sucrose well at times relative to analysis events. Similarly, in Experiment 2, lever presses were recorded to calculate the hazard rate and latency to press after lever extension, and feeder port entries and exits were recorded to calculate feeder entry probability and cumulative feeder entries during auditory cue presentation.

For most comparisons, except where otherwise noted, mean phasic firing was calculated as the mean firing rate in the period 30-500 ms after the event in question minus the average firing rate 0-1000 ms before the baseline event. In Experiment 1, the baseline event was uniformly defined as cue onset for each trial. In Experiment 2, baseline was calculated based on firing rate before lever extension for lever extension and auditory cue, and before light cue for light cue and reward,

In Experiment 2, lever extensions were identical for all three trial types, so responses were aggregated across trial types to construct PSTHs and compute mean responses. For all events following lever press, responses were segregated by trial type (A vs. B vs. C).

Hazard rate was calculated as probability of lever press in a given bin given that no lever press has occurred – i.e. P(LP = t | LP >= t) (Hamid, Pettibone et al. 2016). For each session, time following lever extension was binned in 100 ms intervals, and hazard rate in each bin was calculated as the percent of trials in which lever press occurred in that bin out of all trials in which lever press had not already occurred.

### Histology

After the final recordings, 100 mA current was passed through the recording electrodes to mark their locations. Rats were immediately sacrificed and perfused with cold phosphate buffered saline (PBS) and 4% paraformaldehyde (PFA). Brains were dissected and stored for 24 hours in PFA, followed by PBS. 100 um sections were collected and scanned on a slide-scanning microscope (Olympus) in 4x brightfield. Scans were compared with Paxinos & Watson’s rat brain atlas to determine electrode placement (Supplementary Fig. 10) (Paxinos and Watson 2007).

### Data analysis and statistics

All data analysis was carried out in MATLAB and GraphPad Prism. Residuals were tested for normality, and comparable non-parametric tests were substituted for their canonical parametric counterparts where appropriate. Outliers were detected and removed with Prism’s ROUT method at Q = 1%. The significance level was set at 0.05. * indicates *p* < 0.05, ** *p* < 0.01, *** *p* < 0.001, \*\*\*\**p* < 0.0001.

Specific tests used were as follows: to compare a sample mean to a known reference value: one-sample *t*-test or one-sample Wilcoxon rank test. To compare sample means of two groups with paired observations: two-sample paired *t*-test or Wilcoxon signed-rank test. To compare sample means of two groups without paired observations: two sample unpaired *t*-test or Mann-Whitney *U*-test. To compare more than two groups with matched observations: 1-way repeated measures (RM) ANOVA or Friedman test. To compare more than two groups without matched observations: ordinary 1-way ANOVA or Kruskall-Wallis test. To compare more than two groups with matched observations along two different variables, 2-way RM ANOVA with Fisher’s least significance difference (LSD) test for multiple comparisons, or 2-way RM mixed-effects analysis with Fisher’s LSD test for multiple comparisons. To compare actual to expected categorical counts, Chi-squared test or Fisher’s exact test.

### Data and code availability

All data and code will be available upon reasonable request.

## Acknowledgments

We thank Y. Takahashi for support in building the electrode bundles and microdrives. We thank S. Ahmed, J. Berke, N. Eshel, H. Fields, P. Kalivas, S. Mahler, E. Margolis, and Y. Shaham for comments and suggestions on the general conceptual framework and/or the manuscript.

## Author contributions

Conceptualization: CML, GS, KM

Methodology: CML, NEM, VSN, KMC, GS, KM

Investigation: CML, NEM, VSN, KM

Visualization: CML, NEM, KM

Funding acquisition: GS, KM

Project administration: KM

Supervision: KM

Writing – original draft: CML, KM

Writing – review & editing: CML, NEM, VSN, KMC, GS, KM

## References

Ahmed, S. H., M. Lenoir and K. Guillem (2013). “Neurobiology of addiction versus drug use driven by lack of choice.” Curr Opin Neurobiol 23(4): 581–587.

Azcorra, M., Z. Gaertner, C. Davidson, Q. He, H. Kim, S. Nagappan, C. K. Hayes, C. Ramakrishnan, L. Fenno, Y. S. Kim, K. Deisseroth, R. Longnecker, R. Awatramani and D. A. Dombeck (2023). “Unique functional responses differentially map onto genetic subtypes of dopamine neurons.” Nat Neurosci 26(10): 1762–1774.

Berke, J. D. and S. E. Hyman (2000). “Addiction, dopamine, and the molecular mechanisms of memory.” Neuron 25(3): 515–532.

Berridge, K. C. and T. E. Robinson (2016). “Liking, wanting, and the incentive-sensitization theory of addiction.” Am Psychol 71(8): 670–679.

Caprioli, D., T. Zeric, E. B. Thorndike and M. Venniro (2015). “Persistent palatable food preference in rats with a history of limited and extended access to methamphetamine self-administration.” Addict Biol 20(5): 913–926.

Chow, J. J. and J. S. Beckmann (2021). “Remifentanil-food choice follows predictions of relative subjective value.” Drug Alcohol Depend 218: 108369.

Cohen, J. Y., S. Haesler, L. Vong, B. B. Lowell and N. Uchida (2012). “Neuron-type-specific signals for reward and punishment in the ventral tegmental area.” Nature 482(7383): 85–88.

Crespo, J. A., K. Sturm, A. Saria and G. Zernig (2005). “Simultaneous intra-accumbens remifentanil and dopamine kinetics suggest that neither determines within-session operant responding.” Psychopharmacology (Berl) 183(2): 201–209.

Daw, N. D., S. J. Gershman, B. Seymour, P. Dayan and R. J. Dolan (2011). “Model-based influences on humans’ choices and striatal prediction errors.” Neuron 69(6): 1204–1215.

Di Chiara, G. (1999). “Drug addiction as dopamine-dependent associative learning disorder.” Eur J Pharmacol 375(1-3): 13–30.

Diana, M., A. L. Muntoni, M. Pistis, M. Melis and G. L. Gessa (1999). “Lasting reduction in mesolimbic dopamine neuronal activity after morphine withdrawal.” Eur J Neurosci 11(3): 1037–1041.

Doll, B. B., D. A. Simon and N. D. Daw (2012). “The ubiquity of model-based reinforcement learning.” Curr Opin Neurobiol 22(6): 1075–1081.

Eshel, N., M. Bukwich, V. Rao, V. Hemmelder, J. Tian and N. Uchida (2015). “Arithmetic and local circuitry underlying dopamine prediction errors.” Nature 525(7568): 243–246.

Eshel, N., J. Tian, M. Bukwich and N. Uchida (2016). “Dopamine neurons share common response function for reward prediction error.” Nat Neurosci 19(3): 479–486.

Ferguson, L. M., A. M. Ahrens, L. G. Longyear and J. W. Aldridge (2020). “Neurons of the Ventral Tegmental Area Encode Individual Differences in Motivational “Wanting” for Reward Cues.” J Neurosci 40(46): 8951–8963.

FitzGerald, T. H., R. J. Dolan and K. Friston (2015). “Dopamine, reward learning, and active inference.” Front Comput Neurosci 9: 136.

Gardner, M. P. H., G. Schoenbaum and S. J. Gershman (2018). “Rethinking dopamine as generalized prediction error.” Proc Biol Sci 285(1891).

Gershman, S. J. and N. Uchida (2019). “Believing in dopamine.” Nat Rev Neurosci 20(11): 703–714.

Hamid, A. A., J. R. Pettibone, O. S. Mabrouk, V. L. Hetrick, R. Schmidt, C. M. Vander Weele, R. T. Kennedy, B. J. Aragona and J. D. Berke (2016). “Mesolimbic dopamine signals the value of work.” Nat Neurosci 19(1): 117–126.

Jeong, H., A. Taylor, J. R. Floeder, M. Lohmann, S. Mihalas, B. Wu, M. Zhou, D. A. Burke and V. M. K. Namboodiri (2022). “Mesolimbic dopamine release conveys causal associations.” Science 378(6626): eabq6740.

Kalhan, S., A. D. Redish, R. Hester and M. I. Garrido (2021). “A salience misattribution model for addictive-like behaviors.” Neurosci Biobehav Rev 125: 466–477.

Kalivas, P. W. and N. D. Volkow (2005). “The neural basis of addiction: a pathology of motivation and choice.” Am J Psychiatry 162(8): 1403–1413.

Kalmbach, A., V. Winiger, N. Jeong, A. Asok, C. R. Gallistel, P. D. Balsam and E. H. Simpson (2022). “Dopamine encodes real-time reward availability and transitions between reward availability states on different timescales.” Nat Commun 13(1): 3805.

Keiflin, R. and P. H. Janak (2015). “Dopamine Prediction Errors in Reward Learning and Addiction: From Theory to Neural Circuitry.” Neuron 88(2): 247–263.

Langdon, A. J., M. J. Sharpe, G. Schoenbaum and Y. Niv (2018). “Model-based predictions for dopamine.” Curr Opin Neurobiol 49: 1–7.

Lefevre, E. M., M. T. Pisansky, C. Toddes, F. Baruffaldi, M. Pravetoni, L. Tian, T. J. Y. Kono and P. E. Rothwell (2020). “Interruption of continuous opioid exposure exacerbates drug-evoked adaptations in the mesolimbic dopamine system.” Neuropsychopharmacology 45(11): 1781–1792.

Leyton, M. (2022). “Do stimulant medications produce sensitization in humans?” Neurosci Biobehav Rev 137: 104657.

Mahler, S. V. and H. de Wit (2010). “Cue-reactors: individual differences in cue-induced craving after food or smoking abstinence.” PLoS One 5(11): e15475.

Marks, K. R., D. N. Kearns, C. J. Christensen, A. Silberberg and S. J. Weiss (2010). “Learning that a cocaine reward is smaller than expected: A test of Redish’s computational model of addiction.” Behav Brain Res 212(2): 204–207.

McHugh, R. K., S. Park and R. D. Weiss (2014). “Cue-induced craving in dependence upon prescription opioids and heroin.” Am J Addict 23(5): 453–458.

Mohebi, A., V. L. Collins and J. D. Berke (2023). “Accumbens cholinergic interneurons dynamically promote dopamine release and enable motivation.” Elife 12.

Mohebi, A., J. R. Pettibone, A. A. Hamid, J. T. Wong, L. T. Vinson, T. Patriarchi, L. Tian, R. T. Kennedy and J. D. Berke (2019). “Dissociable dopamine dynamics for learning and motivation.” Nature 570(7759): 65–70.

Moussawi, K., M. M. Ortiz, S. C. Gantz, B. J. Tunstall, R. C. N. Marchette, A. Bonci, G. F. Koob and L. F. Vendruscolo (2020). “Fentanyl vapor self-administration model in mice to study opioid addiction.” Sci Adv 6(32): eabc0413.

Niv, Y., N. D. Daw, D. Joel and P. Dayan (2007). “Tonic dopamine: opportunity costs and the control of response vigor.” Psychopharmacology (Berl) 191(3): 507–520.

O’Sullivan, S. S., A. H. Evans and A. J. Lees (2009). “Dopamine dysregulation syndrome: an overview of its epidemiology, mechanisms and management.” CNS Drugs 23(2): 157–170.

Ostlund, S. B., K. H. LeBlanc, A. R. Kosheleff, K. M. Wassum and N. T. Maidment (2014). “Phasic mesolimbic dopamine signaling encodes the facilitation of incentive motivation produced by repeated cocaine exposure.” Neuropsychopharmacology 39(10): 2441–2449.

Panlilio, L. V. and C. W. Schindler (2000). “Self-administration of remifentanil, an ultra-short acting opioid, under continuous and progressive-ratio schedules of reinforcement in rats.” Psychopharmacology (Berl) 150(1): 61–66.

Panlilio, L. V., E. B. Thorndike and C. W. Schindler (2007). “Blocking of conditioning to a cocaine-paired stimulus: testing the hypothesis that cocaine perpetually produces a signal of larger-than-expected reward.” Pharmacol Biochem Behav 86(4): 774–777.

Redish, A. D. (2004). “Addiction as a computational process gone awry.” Science 306(5703): 1944–1947.

Robbins, T. W. (1978). “The acquisition of responding with conditioned reinforcement: effects of pipradrol, methylphenidate, d-amphetamine, and nomifensine.” Psychopharmacology (Berl) 58(1): 79–87.

Saal, D., Y. Dong, A. Bonci and R. C. Malenka (2003). “Drugs of abuse and stress trigger a common synaptic adaptation in dopamine neurons.” Neuron 37(4): 577–582.

Sadacca, B. F., J. L. Jones and G. Schoenbaum (2016). “Midbrain dopamine neurons compute inferred and cached value prediction errors in a common framework.” Elife 5.

Samaha, A. N., S. Y. Khoo, C. R. Ferrario and T. E. Robinson (2021). “Dopamine ‘ups and downs’ in addiction revisited.” Trends Neurosci 44(7): 516–526.

Saraiya, T. C., A. M. Jarnecke, J. Jones, D. G. Brown, K. T. Brady and S. E. Back (2021). “Laboratory-induced stress and craving predict opioid use during follow-up among individuals with prescription opioid use disorder.” Drug Alcohol Depend 225: 108755.

Schultz, W. (2016). “Dopamine reward prediction-error signalling: a two-component response.” Nat Rev Neurosci 17(3): 183–195.

Schultz, W., P. Dayan and P. R. Montague (1997). “A neural substrate of prediction and reward.” Science 275(5306): 1593–1599.

Sinha, R. (2011). “New findings on biological factors predicting addiction relapse vulnerability.” Curr Psychiatry Rep 13(5): 398–405.

Steinberg, E. E., R. Keiflin, J. R. Boivin, I. B. Witten, K. Deisseroth and P. H. Janak (2013). “A causal link between prediction errors, dopamine neurons and learning.” Nat Neurosci 16(7): 966–973.

Steketee, J. D. and P. W. Kalivas (2011). “Drug wanting: behavioral sensitization and relapse to drug-seeking behavior.” Pharmacol Rev 63(2): 348–365.

Strang, J., N. D. Volkow, L. Degenhardt, M. Hickman, K. Johnson, G. F. Koob, B. D. L. Marshall, M. Tyndall and S. L. Walsh (2020). “Opioid use disorder.” Nat Rev Dis Primers 6(1): 3.

Takahashi, Y. K., T. A. Stalnaker, L. E. Mueller, S. K. Harootonian, A. J. Langdon and G. Schoenbaum (2023). “Dopaminergic prediction errors in the ventral tegmental area reflect a multithreaded predictive model.” Nat Neurosci 26(5): 830–839.

Tan, B., C. J. Browne, T. Nobauer, A. Vaziri, J. M. Friedman and E. J. Nestler (2024). “Drugs of abuse hijack a mesolimbic pathway that processes homeostatic need.” Science 384(6693).

Taylor, J. R. and T. W. Robbins (1984). “Enhanced behavioural control by conditioned reinforcers following microinjections of d-amphetamine into the nucleus accumbens.” Psychopharmacology (Berl) 84(3): 405–412.

Trifilieff, P. and D. Martinez (2014). “Blunted dopamine release as a biomarker for vulnerability for substance use disorders.” Biol Psychiatry 76(1): 4–5.

Volkow, N. D., J. S. Fowler, G. J. Wang and J. M. Swanson (2004). “Dopamine in drug abuse and addiction: results from imaging studies and treatment implications.” Mol Psychiatry 9(6): 557–569.

Volkow, N. D., G. J. Wang, J. S. Fowler, D. Tomasi and F. Telang (2011). “Addiction: beyond dopamine reward circuitry.” Proc Natl Acad Sci U S A 108(37): 15037–15042.

Voon, V., K. Hassan, M. Zurowski, S. Duff-Canning, M. de Souza, S. Fox, A. E. Lang and J. Miyasaki (2006). “Prospective prevalence of pathologic gambling and medication association in Parkinson disease.” Neurology 66(11): 1750–1752.

Wyvell, C. L. and K. C. Berridge (2001). “Incentive sensitization by previous amphetamine exposure: increased cue-triggered “wanting” for sucrose reward.” J Neurosci 21(19): 7831–7840.

Zijlstra, F., J. Booij, W. van den Brink and I. H. Franken (2008). “Striatal dopamine D2 receptor binding and dopamine release during cue-elicited craving in recently abstinent opiate-dependent males.” Eur Neuropsychopharmacol 18(4): 262–270.

## Methods references

Moussawi, K., W. Zhou, H. Shen, C. M. Reichel, R. E. See, D. B. Carr and P. W. Kalivas (2011). “Reversing cocaine-induced synaptic potentiation provides enduring protection from relapse.” Proc Natl Acad Sci U S A 108(1): 385-390.

Paxinos, G. and C. Watson (2007). The rat brain in stereotaxic coordinates. Oxford, Academic Press.

Richardson, N. R. and D. C. Roberts (1996). “Progressive ratio schedules in drug self-administration studies in rats: a method to evaluate reinforcing efficacy.” J Neurosci Methods 66(1): 1-11.

Roesch, M. R., D. J. Calu and G. Schoenbaum (2007). “Dopamine neurons encode the better option in rats deciding between differently delayed or sized rewards.” Nat Neurosci 10(12): 1615-1624.

Takahashi, Y. K., H. M. Batchelor, B. Liu, A. Khanna, M. Morales and G. Schoenbaum (2017). “Dopamine Neurons Respond to Errors in the Prediction of Sensory Features of Expected Rewards.” Neuron 95(6): 1395-1405 e1393.

