## Supplementary Figures/Notes for "Generalized cue reactivity in dopamine neurons after opioids"

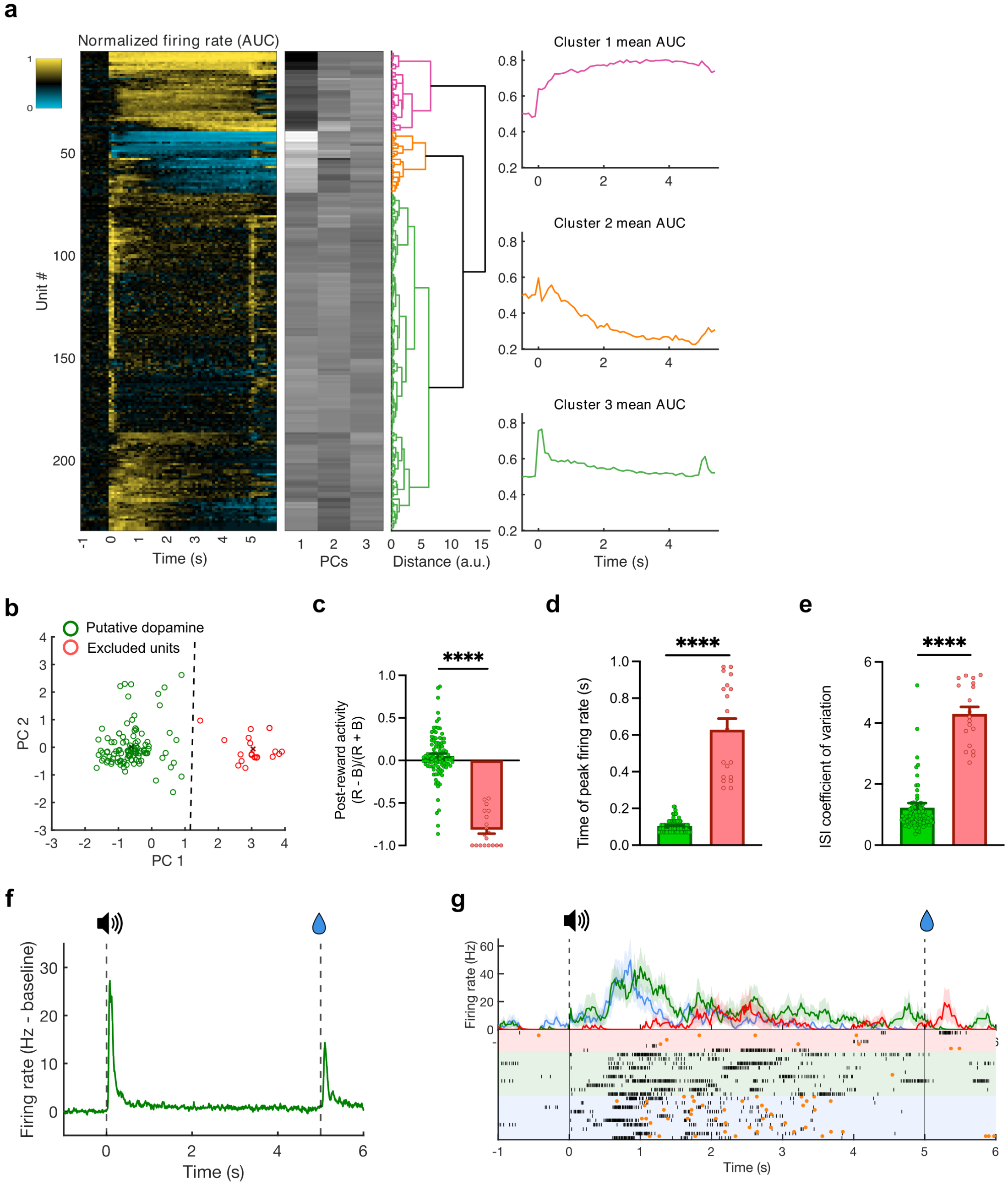


**Supplementary Fig. 1. Putative dopamine neurons identification in Experiment 1.** (**a**) Raw unit clustering results from Experiment 1. Left: heatmap showing functional activation of each unit (rows) aligned to sucrose cue onset (0 s). Activity normalized using area under receiver-operator curve (auROC) method, compared against baseline activity during 5 s before cue onset. Scales from 0 (signal perfectly discriminable, less than baseline) to 1 (perfectly discriminable, greater than baseline). Center left: first three components extracted via PCA. Center: dendrogram showing results from hierarchical agglomerative clustering. Right: mean AUC normalized activity for units classified into each cluster. (**b**) Scatter plot of first two principal components extracted from peak firing 50-1000 ms after cue, peri-reward inhibition, and ISI coefficient of variation for all units in cluster 3 with baseline firing rate <= 12Hz. Green circles putative dopamine units retained by k-means clustering; red circles were eliminated units, with centroids marked by corresponding colored crosses. Black dashed line indicates decision boundary. (**c**) Time of peak firing 50-1000 ms after cue for retained (green) and excluded units (red); Mann-Whitney U-test, $n = 119$, $U = 67.50$, $p < 0.0001$. (**d**) Peri-reward inhibition ([R-B]/[R+B] where R = firing rate 0-1 s after reward, and B = baseline firing rate) for retained and excluded units ($U = 145$, $p < 0.0001$). (**e**) Coefficient of variation of ISIs for spikes recorded outside phasic firing periods (~500 ms after programmed events) for retained and excluded units ($U = 21$, $p < 0.0001$). (**f**) Mean ± SEM firing rate (Hz – baseline) of units from raw phasic firing cluster that were retained as putative dopamine (n = 100). (**g**) Example PSTH and raster session data from one eliminated neuron from the original cluster 3. Vertical lines indicate cue onset (0 s) and cue offset/sucrose delivery (5 s). Trials binned at 20 ms for clarity. All traces (red = neutral trials, green = RMF, and blue = sucrose) and shaded outlines indicate mean ± SEM. Raster plot shows individual trials. Black ticks mark recorded spikes. Orange dots indicate sucrose port entries.

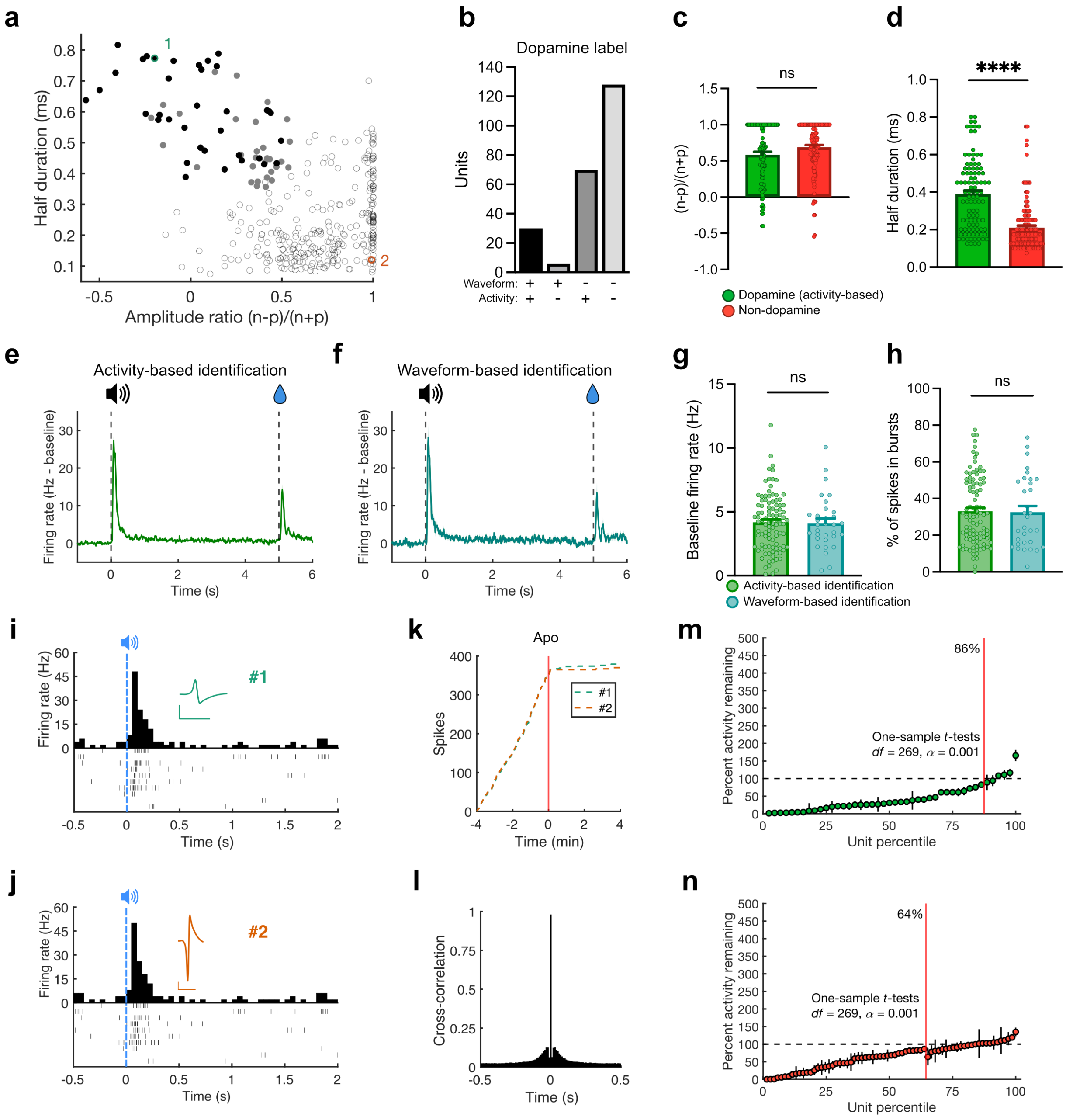


**Supplementary Fig. 2. Waveform-based identification of dopamine neurons was more conservative but consistent with functional clustering.**

(**a**) Scatter plot of all units by half-duration and amplitude ratio. Black-filled circles are identified as dopamine neurons by waveform-based k-means clustering (Roesch, Calu et al. 2007, Takahashi, Batchelor et al. 2017). Gray-filled circles are non-responsive units in the activity-based dopamine cluster from Fig. 2. Empty, semi-opaque circles are classified as non-dopamine. Identified units 1 and 2 were recorded in different channels in the same session and likely represent the same neuron; this is described further in panels e-h. (**b**) Number of units identified as dopamine by waveform criteria (panel a) and activity-based criteria in Fig. 2. (**c**) Amplitude ratios of units identified as dopamine by activity-based method (Mann-Whitney U-test, $n = 234$, $U = 5920$, $p = 0.1304$). (**d**) Half-durations of units identified as dopamine by activity-based method (Mann-Whitney U-test, $n = 234$, $U = 2791$, $p < 0.0001$). (**e**) Trace of mean ± SEM firing response during sucrose trials for units identified by activity-based clustering ($n$ = 99). (**f**) Trace of mean ± SEM firing response during sucrose trials for units identified by waveform criteria ($n$ = 32). (**g**) Baseline firing rates for neurons identified as dopamine based on sucrose trial spiking activity (green) and waveform properties (teal; Mann-Whitney $U$ test, $n$ = 131, $U$ = 1531, $p$ = 0.7765). (**h**) Percent of recorded spikes occurring in bursts for neurons identified as dopamine based on sucrose trial spiking activity and waveform properties (Mann-Whitney $U$ test, $n$ = 131, $U$ = 1536, $p$ = 0.7970). (**i**) Raster plots and PSTH of example responses to sucrose cue from Unit 1 in panel A, identified as dopaminergic by waveform properties. Waveform inset at scale of 500 ms, 75 µV. (**j**) Raster plots and PSTH from Unit 2 in panel A identified as non-dopaminergic, during the same behavioral trials as in panel I (both units recorded in different channels during the same session). Note that firing of the 2 units appears identical in response to the sucrose cue. (**k**) Cumulative spikes around IV infusion of 20 µg/kg apomorphine for units 1 and 2. (**l**) Cross-correlation of spikes in units 1 and 2. Note correlation is approximately 1 at zero delay suggesting the two recordings represent the same neuron. (**m**) 86% of dopamine neurons show inhibition of firing after apomorphine infusion. Neuronal firing relative to baseline (mean ± 95% CI) after IV apomorphine infusion in activity-identified dopamine neurons (baseline: five min prior to infusion; mean activity: 5 min after infusion). For each individual neuron, firing in the interval 30-300 s post-infusion was divided into 1 s intervals, and tested with a one-sample t-test for difference from pre-infusion mean baseline firing rate at alpha = 0.001. Vertical bars indicate 95% confidence interval of inhibition, with large bars suggesting greater variability of post-infusion firing rate. Horizontal black dashed line indicates line of no-effect. Vertical red line separates units with significant from non-significant response inhibition by apomorphine. (**n**) Same as panel m but for activity-identified non-dopamine neurons.

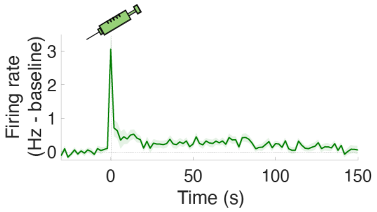


**Supplementary Fig 3. Persistent dopamine neuron response to 4 µg/kg remifentanil**
Trace (mean ± SEM) of firing responses to cued RMF infusion of 4 µg/kg across recordings (*n* =31).

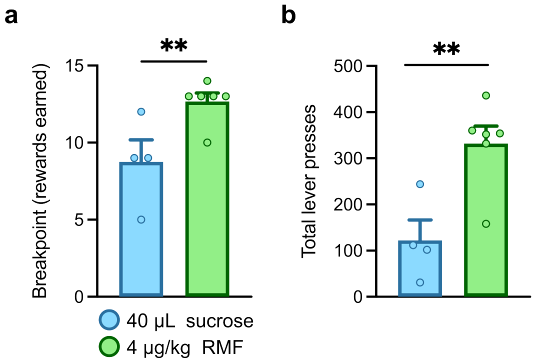


**Supplementary Fig 4. Breakpoint for RMF and sucrose in a progressive ratio schedule of reinforcement.**
(**a**) Training was conducted at the end of Experiment 1 in the opioid-exposed group. Final step achieved during progressive ratio (PR) test in which lever press requirement to receive reward (40 µL sucrose or 4 µg/kg RMF) was increased according to an exponential schedule (Richardson and Roberts 1996) (steps = 1, 2, 4, 6, 9, 12, 15, 20, 25, 32, 40, 50, 62, 77, 95, 118, 145, 178, 219, 268, 328, 402, 492, 603, and 737). Each point represents the highest level achieved out of ~3 PR attempts per rat (Mann-Whitney U-test, n = 10, U = 1, p = 0.0095). Note that initially 6 rats were trained to respond for RMF. One rat failed to acquire lever pressing behavior for sucrose reward, and one rat was sacrificed before sucrose training could be completed. (**b**) Total lever presses on active lever during best PR performances for sucrose and RMF reward per rat (Unpaired *t*-test, $t(8) = 3.566$, $p = 0.0073$).

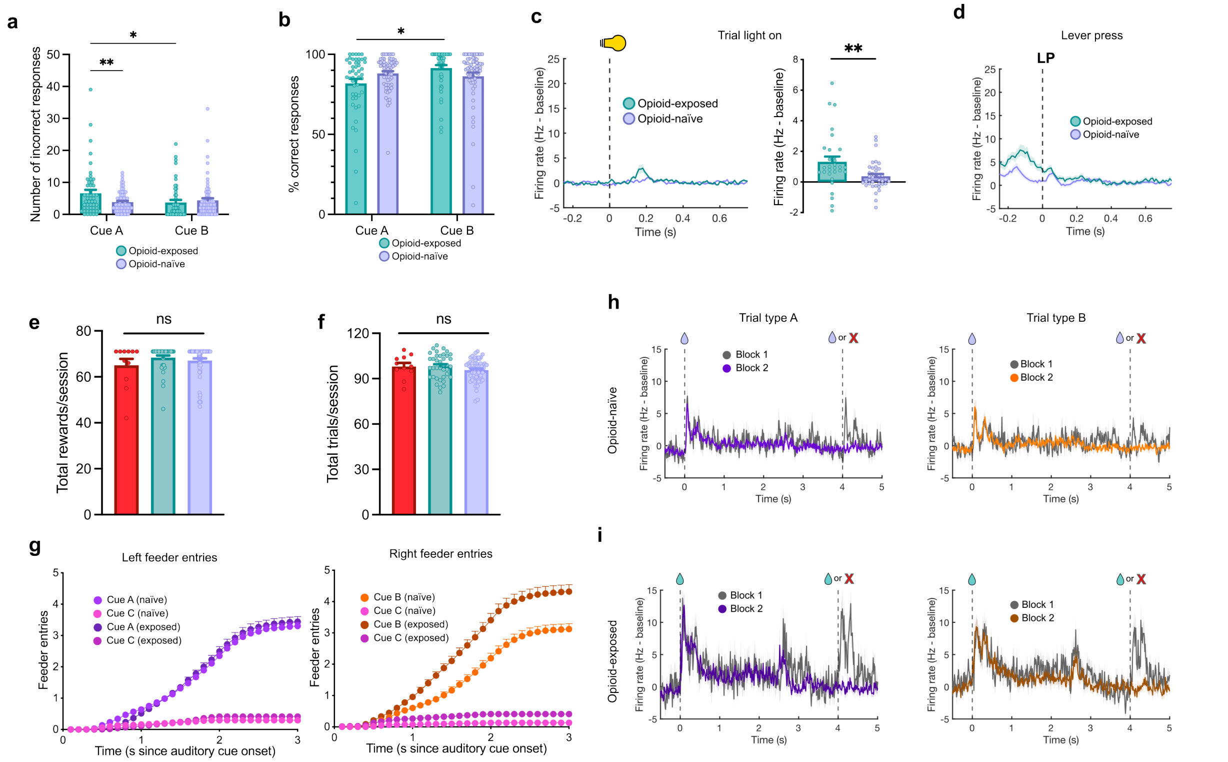


**Supplementary Fig 5. Behavioral performance and dopamine firing response data from the operant procedure.**

(**a**) Number of incorrect responses per recording session to Cue A and Cue B for opioid-exposed and opioid-naïve rats (2-way RM ANOVA, Cue x Exposure interaction: $F(1,114) = 4.476$, $p = 0.0366$). (**b**) Percent correct feeder-in responses to auditory Cue A and Cue B per recording session in opioid-exposed and opioid-naïve rats (Cue factor: $F(1,114) = 3.245$, $p = 0.0743$; Exposure factor: $F(1,114) = 0.5252$, $p = 0.9907$; Cue x Exposure interaction $F(1,114) = 3.098$, $p = 0.0811$). (**c**) Left: Trace of average putative dopamine responses to trial light illumination at the start of trial for opioid-exposed and opioid-naïve units, prior to lever extension. Right: average firing response during 100-200 ms after trial light turns on (Mann-Whitney test, $n = 68$, $U = 330$, $p = 0.0027$). (**d**) Trace of average putative dopamine response to lever press for opioid-exposed and opioid-naïve units. (**e**) Total number of rewards during each session for rat 8 (red) all other opioid-exposed rats (teal) and opioid-naïve rats (purple; Kruskal-Wallis test, $n = 107$, $k = 2.328$, $p = 0.3122$). (**f**) Total number of trials (all types) during each session for rat 8 (red) all other opioid-exposed rats (teal) and opioid-naïve rats (purple Kruskal-Wallis test, $n = 107$, $k = 2.338$, $p = 0.3107$). (**g**) Cumulative feeder entries at the non-drug (left panel) and drug-associated feeder (right panel) following auditory cue onset. Note low rate of responding to Cue C. (**h**) Average responses of opioid-naïve units to reward delivery in block 1 (gray) and block 2 (color), with reward one delivered at 0 s and reward two (or omission) at 4 s. Left panel shows responses to reward at left port in trial-type A (purple/gray) and right panel shows reward at right port in trial-type B (orange/gray). (**i**) Same as in G but for opioid-exposed units. Note the absence of negative RPEs in either group to the second reward omission, likely because the animals are well trained and learned to expect the second reward omission.

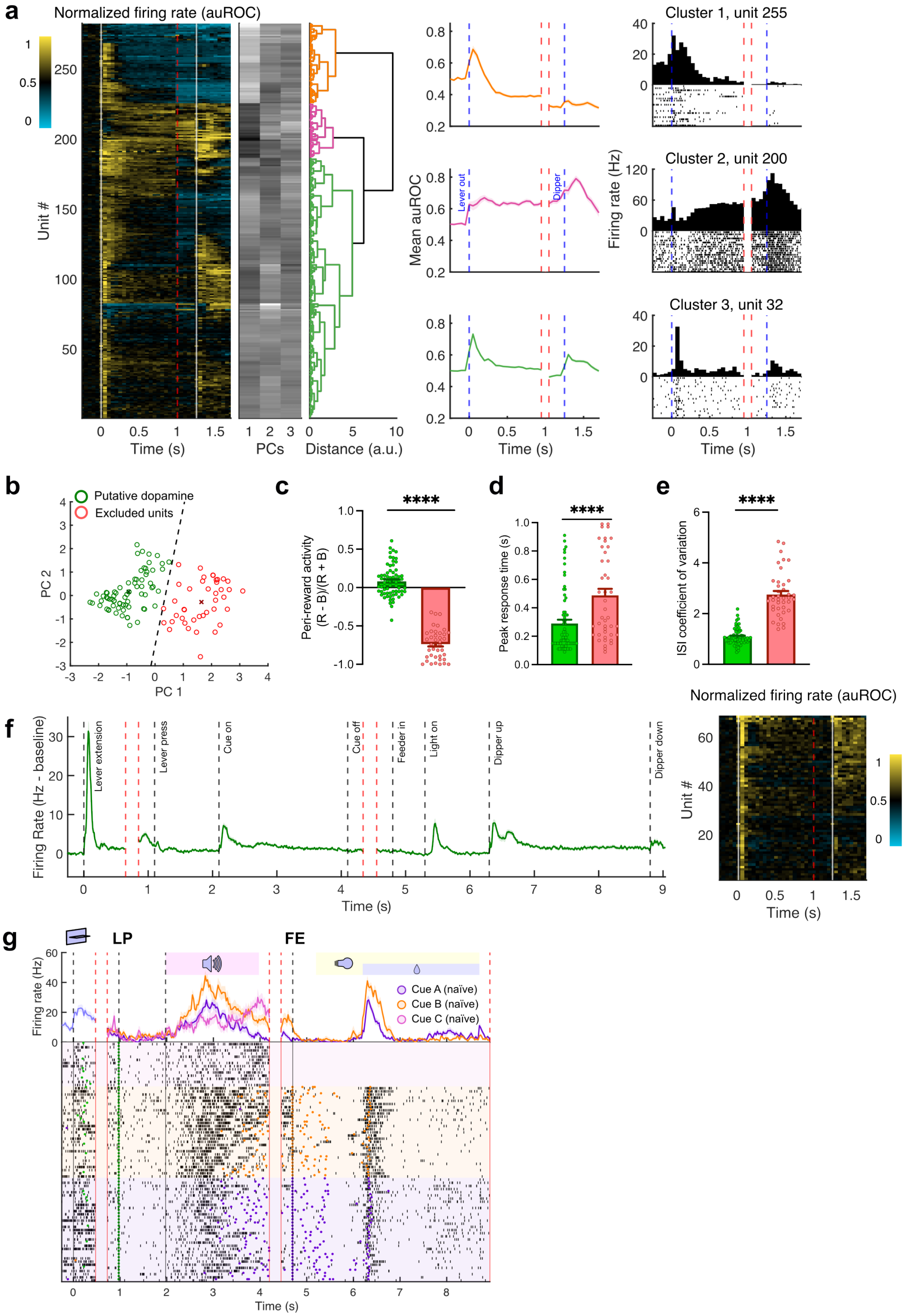


**Supplementary Fig 6. Putative dopamine neurons identification in Experiment 2.**

(**a**) Initial clustering results for cue-responsive units in operant experiment. Left: Heatmap showing functional activation of each unit (rows) centered on lever extension and water reward delivery. Events are displayed as white solid lines, and time discontinuity as dashed red line. Activity normalized using area under receiver-operator curve (auROC) method, compared against baseline. Scales from 0 to 1. Center left, first three components extracted via PCA. Center: Dendrogram showing results from hierarchical clustering split at three clusters. Center right: Mean ± SEM auROC data for each group. Time discontinuity shown as white space flanked by dashed red lines. Right: example neurons from each cluster. PSTH and raster plots from example units for each cluster. (**b**) Scatter plot of first two PCs extracted from peak firing 50-1000 ms after cue, peri-reward inhibition, and ISI coefficient of variation for all units in cluster with baseline firing rate <= 12Hz. Green circles putative dopamine units retained by k-means clustering; red circles were eliminated units, with centroids marked by corresponding colored crosses. Black dashed line indicates decision boundary. (**c**) Peri-reward inhibition ([R-B]/[R+B] where R = firing rate 0-1 s either before or after reward, and B = baseline firing rate) for retained putative dopamine (green) and excluded units (red; Mann-Whitney U-test, $n = 109$, $U = 3$, $p < 0.0001$). (**d**) Time of peak firing 50-1000 ms after auditory cue for retained and excluded units ($n = 109$, $U = 726.5$, $p < 0.0001$). (**e**) Coefficient of variation of ISIs for spikes recorded outside phasic firing periods (~500 ms after programmed events) for retained (green) and excluded units (red; Mann-Whitney U-test, $n = 109$ $U = 45$, $p < 0.0001$). (**f**) Left; Mean ± SEM firing rate responses for neurons retained as putative dopamine neurons (n = 69). Right; corresponding auROC heatmap of final putative dopamine cluster. (**g**) Example PSTH and raster session data from one eliminated neuron from the original cluster 3. Purple and orange dots indicate left and right feeder entries respectively. Green dots indicate lever presses.

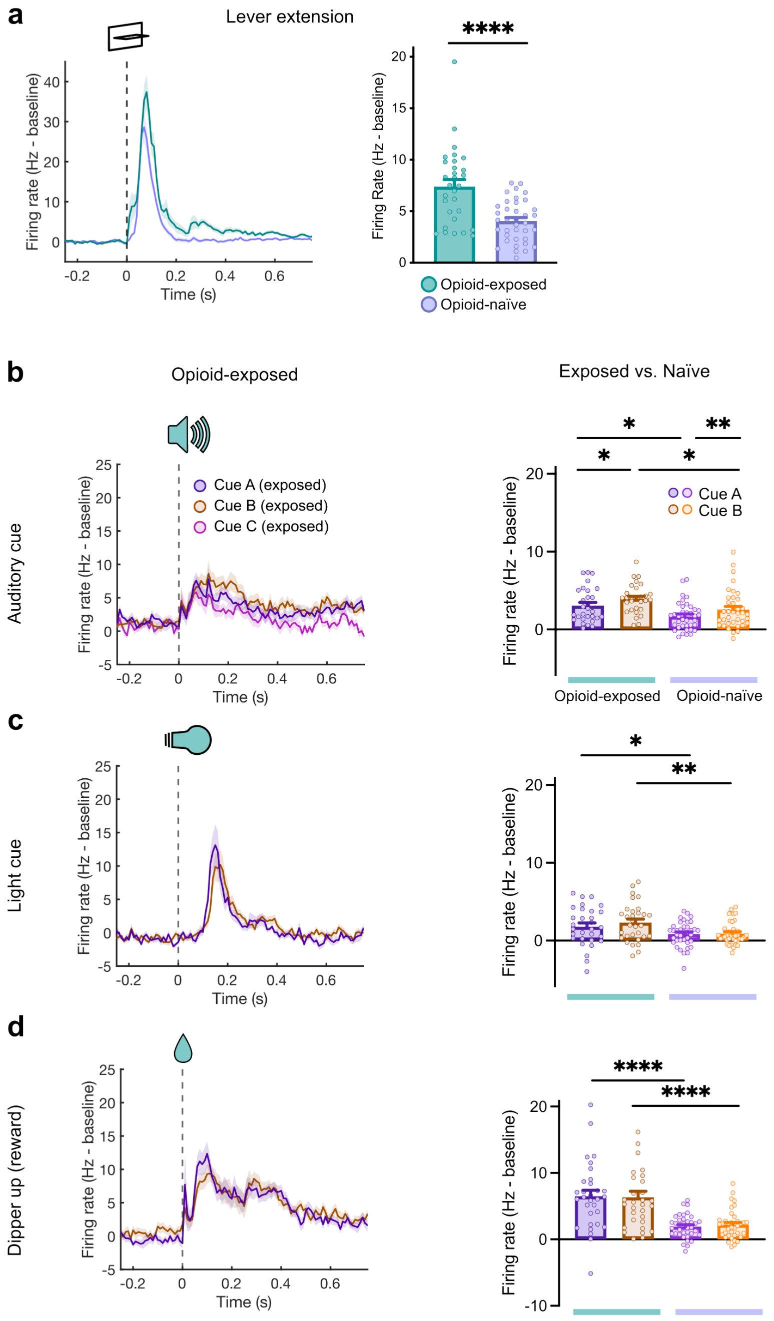


**Supplementary Fig 7. Sensitized dopamine responses to drug and non-drug cues in all opioid-exposed rats.**

Dopamine firing response analyses comparing opioid-naïve to opioid-exposed units including those observed in rat 8. Panels are similar to those in Fig. 5. (**a**) Left: PSTH of baseline-subtracted firing rate around lever extension for opioid-exposed and opioid-naïve units. Right: Mean firing rate for both groups (Mann-Whitney test, $n = 67$, $U = 242$, $p < 0.0001$). (**b**) PSTH of the responses for opioid-exposed units around to auditory cues. Mean firing rates for opioid-exposed and opioid-naïve units (same naïve units as in Fig. 5 g) (2-way RM mixed-effects analysis, Cue factor: $F(1,64) = 10.79$, $p = 0.0017$; Exposure factor: $F(1,65) = 8.157$, $p = 0.0058$; Cue x Exposure interaction $F(1,64) = 0.04719$, $p = 0.8287$). (**c**) Responses of opioid-exposed and opioid-naïve neurons to light cue, similar to b. Comparison of mean baseline-subtracted firing rates light cues in both groups (Cue factor: $F(1,65) = 1.321$, $p = 0.2547$; Exposure factor: $F(1,67) = 7.537$, $p = 0.0078$; Cue x Exposure interaction $F(1,65) = 1.541$, $p = 0.2190$. (**d**) Responses of opioid-exposed and opioid-naïve neurons to water reward delivery (dipper up). Comparison of mean baseline-subtracted firing rate after left vs. right dipper up for opioid-exposed and opioid-naïve units (Cue factor: $F(1,66) = 0.2852$, $p = 0.5951$; Exposure factor: $F(1,67) = 24.37$, $p < 0.0001$; Cue x Exposure interaction $F(1,66) = 0.003168$, $p = 0.9553$).

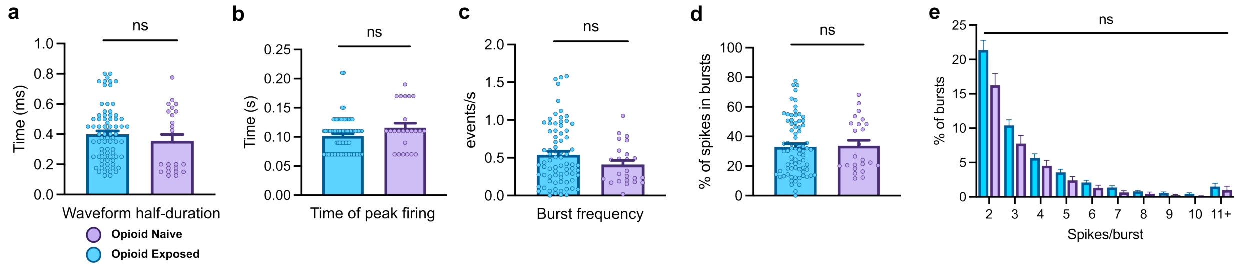


**Supplementary Fig 8. Dopamine neurons waveform and firing properties between groups.** (**a**) Waveform half-duration of putative dopamine neurons for opioid-exposed vs. opioid-naïve units from Experiment 1, defined as time from negative peak to next positive peak, or end of waveform (Mann-Whitney test, $n = 99$, $U = 777$, $p = 0.3176$). (**b**) Time of peak firing during first second following sucrose cue onset ($n = 99$, $U = 822$, $p = 0.5245$). (**c**) Frequency of bursts during ITI for opioid-exposed and naïve units ($n = 99$, $U = 754$, $p = 0.2357$). (**d**) Percent of all recorded ITI spikes occurring in bursts ($n = 99$, $U = 867$, $p = 0.7921$). (**e**) Percentages of detected bursts during ITIs containing $n$ spikes (Two-way ANOVA, opioid effect: $F(1,96) = 2.962$, $p = 0.0885$).

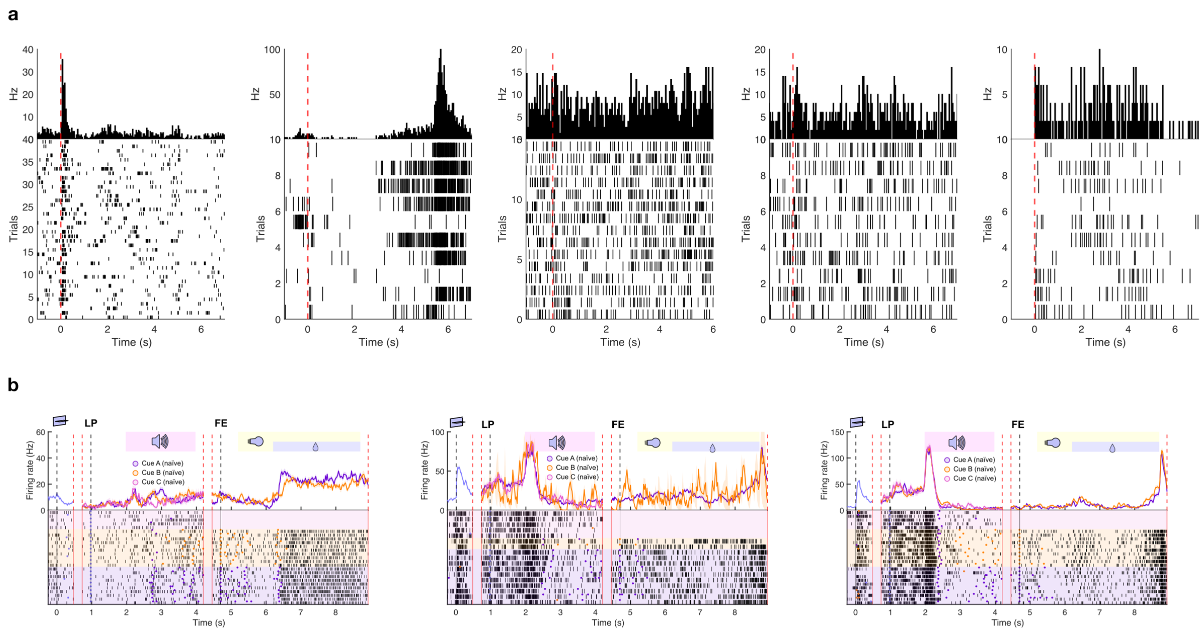

**Supplementary Fig 9. Excluded units from Experiments 1 and 2.**
(**a**), PSTH and raster data from the five units manually removed from the putative dopamine cluster in Experiment 1. Activity centered around sucrose cue onset (0 s). The first unit on the upper left was removed to avoid duplication as it was simultaneously recorded in a nearby channel (Supplementary Fig. 2, i to l). The other units were removed due to concern of misclassification (lack of phasic cue response). (**b**), PSTH and raster data from the three units manually removed from the putative dopamine cluster in Experiment 2, displayed as in Fig. 5. The units were removed due to concern of misclassification (sustained non-phasic responses in unit 1 and extreme outliers in firing response to auditory cue in units 2 and 3).

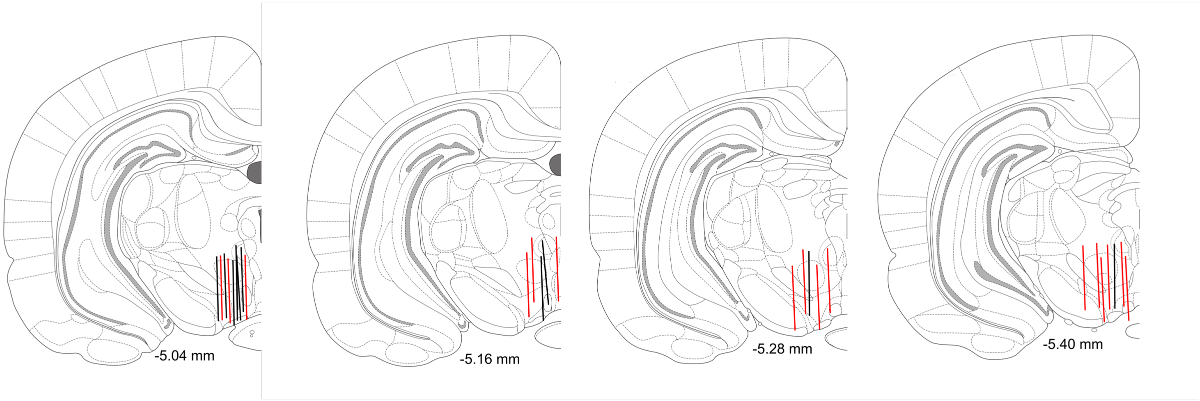


**Supplementary Fig 10. Histological verification of electrode placement.**
Lines show final placement of electrode track. Red lines correspond to electrodes in Experiment 1, and black lines correspond to Experiment 2. Numbers indicate antero-posterior position relative to Bregma.

**Supplementary Notes**

Supplementary Note 1

Data show that blocking effect – the effect of previous learning about a cue to prevent new learning about another cue – continues to occur with addictive drugs (Panlilio, Thorndike et al. 2007), unlike what is predicted by the noncompensable RPE hypothesis. Also, smaller-than-expected drug rewards result in reduced lever pressing suggesting that negative PE could occur for drugs (Marks, Kearns et al. 2010). Although the RPE-based theory's parsimony derives from its adherence to the temporal difference (TD) RPE model of dopamine, this model itself has been criticized, as new evidence has shown that dopamine signals reflect computations that incorporate a richer set of information than the traditional TD account allows (Langdon, Sharpe et al. 2018). Accordingly, several TD-variants (Gershman and Uchida 2019, Takahashi, Stalnaker et al. 2023) and non-TD alternative models (FitzGerald, Dolan et al. 2015, Jeong, Taylor et al. 2022) have been proposed some of which complicate or entirely contradict this RPE model of addiction.

Supplementary Note 2

Hierarchical clustering identifies canonical dopamine neurons based on their firing response to cues and rewards. More traditional dopamine identification methods based on waveform criteria (e.g., initial waveform positivity and waveform duration) and inhibition by a D2 agonist are effective at screening out non-dopaminergic neurons but have been shown to be overly conservative or non-specific (Cohen, Haesler et al. 2012, Mohebi, Pettibone et al. 2019). For example, many dopamine neurons do not express D2 receptors and some genetically identified non-dopamine neurons are inhibited by D2 agonists (Margolis, Lock et al. 2006). Dopamine neurons identified with the activity-based clustering showed longer duration and higher probability of D2 inhibition than non-dopamine neurons. Similarly, K-means clustering based on waveform properties (amplitude ratio and half-duration) (55) yielded concordant but more conservative identification of dopamine neurons (Fig. S2). However, many likely dopamine neurons were excluded with the waveform-based clustering. One neuron was captured in two channels (nearly one-to-one correspondence of spikes between channels) and was classified as likely dopamine neuron based on waveform criteria in one channel, but non-dopamine in the other (Fig. S2). Such differences in waveform properties are likely due to the placement of the recording wires relative to the neuron. Hierarchical activity-based clustering avoids this problem.

Supplementary Note 3

Although Cue C signaled the end of the trial and absence of reward, it resulted in a positive dopamine response comparable to rewarded cues in both opioid-exposed and naïve groups, despite evidence of behavioral discrimination of the cue's identity (Fig. S5G). The response to Cue B was also slightly elevated relative to Cue A in both groups (Fig. 5G). This is likely due to the sensory qualities of the cue itself, as the reward value was identical for both trial types in the opioid-naïve group, which did not display a behavioral preference for Cue B.

Supplementary Note 4

The first PC (PC1) was extracted from the cue reactivity and dopamine response analyses in Fig. 4P and 5K, respectively. We treated the PC1 from the respective analyses as a compounded approximate measure of behavioral cue reactivity and enhanced dopamine responsiveness: In both analyses, PCs 1 and 2 combined accounted for ~85% of the data's variance, of which >50% was captured by PC1 alone. Further, in the behavioral analysis, all included variables were designed to correlate positively with reactivity and loaded positively on PC1. Finally, in both analyses, the opioid-exposed group was largely higher in PC1 than the opioid-naïve group, consistent with our direct comparisons of the underlying individual variables.

Supplementary Note 5

Although we observed a general sensitization across different cue types, this sensitization was not entirely uniform. For instance, the dopamine firing response remained graded by trial type (rewarded vs. unrewarded). This was observed in the delayed 'valuation' phase of the dopamine response to neutral cues and sucrose omission trials where the time course of the dopamine response differed between rewarded and unrewarded trials. Schultz et al. proposed a model for phasic dopamine response in which two components signal salience, then reward value.(Schultz 2016) In the data from our Pavlovian study, RMF, sucrose, and neutral cues all showed a similar positive dopamine firing response during the initial salience-detection period (30-180 ms) in opioid-exposed rats (but not in opioid-naïve rats), suggesting a state of general hypervigilance, at least towards the sensory modality of reward predictive cues. However, during the valuation period (180-500 ms), responses to the neutral cue rapidly dropped, while firing rates stayed above baseline for both RMF and sucrose cues. A similar detection/valuation firing discrimination was observed in sucrose omission trials: a detection phase immediately after cue offset showed greater firing response in opioid-exposed compared to opioid-naïve; however, a valuation phase around expected reward delivery time showed equivalent negative RPEs between groups. Thus, despite their increased excitability, these neurons are still able to signal value-relevant disappointment, as both neutral cue and reward omission predict a longer delay to the next reward. In the operant experiment, there is also evidence for distinct detection/valuation phases in the response to the auditory cue, which is graded according to reward value. Interestingly, however, the detection phase of this cue response is not enhanced in the opioid-exposed group relative to opioid naïve, unlike every other cue examined within the same subjects in the same experiment. One possibility could be that the auditory cue was discriminative in nature and hence possibly involved different regulatory processes of dopamine firing (e.g., attentional top-down control). Another possibility is that opioid-exposed animals experience a steeper temporal discounting factor, resulting in greatly magnified response of cues near reward (light cue) and less enhancement earlier on (auditory cue). However, the enhancement of the dopaminergic response to the lever extension at the beginning of each trial goes against an explanation in terms of harsher discounting.

Supplementary Note 6

Clinical and pre-clinical imaging studies of various substance use disorders show increased mesolimbic dopamine release in response to drugs or drug-associated cues, indicative of a sensitized hyperdopaminergic state (Diana, Muntoni et al. 1999, Zijlstra, Booij et al. 2008, Berridge and Robinson 2016, Lefevre, Pisansky et al. 2020, Samaha, Khoo et al. 2021, Leyton 2022), but other studies suggest that striatal dopamine transmission is blunted in drug-experienced subjects (Volkow, Wang et al. 2011, Trifilieff and Martinez 2014). These disparate findings can be accounted for by considering the time interval since the last drug exposure, context of drug administration or cue presentation, and drug exposure procedure (intermittent vs. continuous) (Lefevre, Pisansky et al. 2020, Samaha, Khoo et al. 2021, Leyton 2022). Because imaging often occurs in a clinical context which is not associated with drug availability and at a time of maximal tolerance, these results may reflect a tolerant state of the dopaminergic system rather than a more dominant dopaminergic sensitization state in drug-associated context (where relapse is most likely) that is most pronounced during drug use or in early abstinence. Alternatively, dopamine release can be heavily influenced by downstream synaptic factors like inputs from cholinergic interneurons onto dopamine axon terminals in the striatum, so striatal dopamine levels may differ substantially from somatic VTA dopamine firing rate (Mohebi, Collins et al. 2023). Hence, the hyperdopaminergic state may be more reflective of the state of the "addicted brain." A further difficulty that arises in comparing these data to the clinical cue reactivity literature is that many cue reactivity procedures reference their results against neutral stimuli, and do not directly compare drug stimuli to other motivationally relevant natural-reward related stimuli (Versace, Engelmann et al. 2017). Additionally, such studies have rarely been carried out alongside neuroimaging in opioid-use disorder.

Supplementary Note 7

In free-choice tasks, animals often exhibit strong (but dose-dependent (Chow and Beckmann 2021) preferences for even modest food rewards over drug rewards (Ahmed, Lenoir et al. 2013, Caprioli, Zeric et al. 2015). On the RPE model, this is difficult to explain, as the runaway overvaluation of the drug state ought in the long run to overwhelm any other state, no matter how rewarding, resulting in almost exclusive preference for drug. If, however, opioid use resulted in augmentation of dopamine activity that drives valuation of all reward cues (as our data show), then an impulsive rat faced with the choice of immediate food reward and pharmacokinetically delayed drug reward could easily choose the former.
